# Phylogenomics resolves major relationships and reveals significant diversification rate shifts in the evolution of silk moths and relatives

**DOI:** 10.1101/517995

**Authors:** CA Hamilton, RA St Laurent, K Dexter, IJ Kitching, JW Breinholt, A Zwick, MJTN Timmermans, JR Barber, AY Kawahara

## Abstract

**Background:** The silkmoths and their relatives constitute the ecologically and taxonomically diverse superfamily Bombycoidea, which includes some of the most charismatic species of Lepidoptera. Despite displaying some of the most spectacular forms and ecological traits among insects, relatively little attention has been given to understanding their evolution and the drivers of their diversity.

**Results:** To begin to address this problem, we created a new Bombycoidea-specific Anchored Hybrid Enrichment (AHE) probe set and sampled up to 571 loci for 117 taxa across all major lineages of the Bombycoidea, producing a well-supported phylogeny. The tree was overall consistent with prior morphological and molecular studies, although some taxa (e.g., *Arotros* Schaus) were misplaced in the Bombycidae and here formally transferred to Apatelodidae. We identified important evolutionary patterns (e.g., morphology, biogeography, and differences in speciation and extinction), and our analysis of diversification rates highlights the stark increases that exist within the Sphingidae (hawkmoths) and Saturniidae (wild silkmoths).

**Conclusions:** We postulate that these rate shifts are due to the well-documented bat-moth “arms race” and differences in selective pressures from insectivorous bats. The study establishes a backbone for future evolutionary, comparative, and taxonomic studies, and presents a modified DNA extraction protocol that allows Lepidoptera specimens to be readily sequenced from pinned natural history collections, succeeding in samples up to 30 years old. Our research highlights the flexibility of AHE to generate genomic data from a wide range of museum specimens, both age and preservation method, and will allow researchers to tap into the wealth of biological data residing in natural history collections around the globe.

## Background

The Bombycoidea include some of the most charismatic moths among all Lepidoptera. This ecologically diverse superfamily comprises ten families, 520 genera, and 6,092 species [1]. Although widespread globally, the highest diversity of bombycoids occurs in the tropics and subtropics. This diversity includes the most spectacular forms (i.e., range of body sizes and wing shapes) and functions (i.e., mimicry, predator avoidance, flight capabilities, and feeding strategies) in the Lepidoptera [2].

A number of bombycoids have become important contributors to human culture, originally as economically important species for sericulture or as agricultural pests, but more recently as model organisms for comparative studies of genetics, development, and physiology [2]. Additionally, many lineages play important roles as pollinators ([3], [4], [5], [6], [7], [8], [9]), or as indicators in biodiversity and habitat quality assessments [10]. Of the 10 families, three contain species that have been used as model organisms (Bombycidae, Saturniidae, and Sphingidae). Unfortunately, relationships among bombycoid families, and especially these three families, have remained largely elusive. For example, Bombycidae have been considered the sister lineage to either Saturniidae or Sphingidae ([11], [12], [13], [14], [15], [16], [17], [18]); but see Breinholt and Kawahara [19]. Overall, relatively little attention has been given to the group with regards to understanding their evolution.

Despite their charisma and intrigue, the lack of a robust phylogeny based on broad and dense taxon sampling across the Bombycoidea is dramatically affecting our ability to answer fundamental questions about the drivers of their diversity. Monophyly of the Bombycoidea has been supported by six morphological synapomorphies [15], but Zwick [20] determined that only two of these were systematically informative: one poorly understood thoracic character [21], and one relating to the arrangement of the forewing vein (A. Zwick, unpublished). Recent molecular studies of bombycoid systematics ([11], [13], [20]) have resulted in substantial differences in terms of relationships from morphology-based phylogenetic hypotheses ([15], [21], [22]). To date, nearly all molecular studies of bombycoids have included fewer than 20 protein-coding genes for ≤50 species (e.g., [11], [13], [23], [24]). Although these studies agreed on the monophyly of the superfamily, many relationships among families, subfamilies, and tribes remained unclear and were characterized by weak branch support or conflicting signal. Modern phylogenomics (e.g., based on Anchored Hybrid Enrichment) has been shown to be effective to resolve relationships among Lepidoptera at multiple taxonomic levels ([16], [25], [26], [27], [28]) including the Bombycoidea ([16], [29]). However, those studies that utilize phylogenomics to resolve inter- and intra-familial relationships within the Bombycoidea have been limited by taxon sampling.

To better answer questions regarding bombycoid evolution, we applied Anchored Hybrid Enrichment (AHE) targeted-sequencing phylogenomics [30]. We developed a new, bombycoid-specific, AHE probe set (here called “BOM1”), redesigned from the “LEP1” probe set of Breinholt et al. [16]. Our new probe set captures “legacy” Sanger sequencing-based loci that were part of existing bombycoid molecular datasets ([11], [13], [24], [31]), enabling the merging of older published datasets with those generated from the BOM1 probe set. To further improve the ability to generate a dataset with greater taxon and locus sampling, we developed a new method for extracting DNA from pinned natural history specimens, and show that our extraction approach is successful in obtaining DNA sequence data for phylogenomics. In total, our dataset resulted in 571 loci for 117 species across Bombycoidea.

We also use this opportunity to utilize the phylogeny to examine patterns of diversification in the superfamily. Bombycoidea are well-known to have multiple different ecological life-history strategies, especially for the Saturniidae and Sphingidae – two lineages that harbor the majority of described bombycoid species ([29], [32], [33], [34]). The divergent life-history strategies of these two families ([35], [36]) has likely played a major role in driving their diversity. For example, the majority of hawkmoths feed as adults, seeking out nectar resources during their relatively long lives (weeks to months). During this time, females experience multiple mating events and retain the eggs internally for long periods to allow egg maturation and host plant discovery [35]. This ecological strategy is significantly different from saturniids ([23], [19], [37]), which depend entirely upon the resources acquired during the larval period. Adult saturniids possess reduced or non-functional mouthparts and lay eggs almost immediately after mating. Furthermore, these lineages possess a number of different traits that appear to be anti-bat adaptations in response to echolocating bats, a lineage thought to have arisen approximately 60 million years ago ([38], [39], [40], [41], [42]). Hawkmoths, especially those in the Sphinginae and Macroglossinae, are strong fliers thought to have evolved hearing organs and ultrasound producing organs capable of jamming bat sonar ([33], [43]). Saturniids, in contrast, lack hearing organs and exhibit erratic evasive flight [35] and hindwing tails that deflect bat echoes ([29], [34]). These traits firmly establish the relative ecological roles of Saturniidae and Sphingidae as an important natural experiment from which we can gather valuable information regarding the evolution of predator/prey interactions in moths and their ultimate effects on the diversification process in general.

## Methods

### The “BOM1” AHE probe set design

Anchored Hybrid Enrichment (AHE) is a targeted-sequencing methodology designed to capture hundreds of unique orthologous loci (i.e., single copy, phylogenetically-informative markers) from across the genome, for resolving both shallow and deep-level evolutionary relationships ([11], [13]). Probes are designed to anchor in conserved regions that are flanked by variable regions randomly spread throughout the genome. This approach creates a diverse set of informative loci that include exons, introns, intergenic, and conserved regions of the genome. Targeted-sequencing approaches, like AHE and UCE (Ultraconserved Elements), provide mechanisms whereby different researchers can confidently and effectively use the same loci for independent projects, allowing for the combination of data across studies.

Breinholt et al. [16] constructed a Lepidoptera Agilent Custom SureSelect Target Enrichment “LEP1” probe kit, designed for 855 loci. However, this probe set is not specific to Bombycoidea, and does not include some of the traditional loci that have been used in phylogenetics of Bombycoidea. In order to build a more Bombycoidea-specific AHE probe set and phylogenomic dataset, we began by modifying the LEP1 kit, evaluating which loci were most phylogenetically-informative within the superfamily, optimizing the set of probes to recover these loci, and including 24 previously-sequenced Sanger-sequenced loci, such as e.g., CO1, CAD, DDC, period, wingless and others from [11], [13], [24], and [31], as well as eight vision-related genes (see Supp. Table 1 for locus names). The probes for these vision-related genes are based phototransduction genes mined from eye or head transcriptomes (unpublished; generated by AYK), and are included for future analyses to investigate their evolution across the superfamily.

To determine the informative loci for BOM1, phylogenetically-informative loci were identified by examining the sequence variation of 56 bombycoid species from across the taxonomic breadth of the superfamily (55 AHE samples, from either Breinholt et al. [16] or generated for this study, plus loci mined from the *Bombyx mori* reference genome [44]). These samples were sequenced and processed using the LEP1 probe kit and Breinholt et al. [16] bioinformatics pipeline [45]. Individual gene trees were generated for each of the 855 loci, using Maximum Likelihood in RAxML v8.2 [46], under a GTRGAMMA model of evolution with 100 non-parametric bootstraps for node support. For each tree, phylogenetic informativeness was calculated using PhyDesign ([47], [48], [49]). Parsimony informative characters and the number of segregating sites were calculated using the R packages ‘phyloch’ [50] and ‘ape’ [51] respectively. Additionally, each phylogeny was scrutinized visually to determine whether branch lengths and topological patterns appeared realistic (i.e., no significant outliers present). Loci deemed to be phylogenetically uninformative, or those that were capturing poorly (<60% of sampled species represented for a locus), were excluded from the probe set. The final BOM1 probe kit comprised 571 loci, 539 of which came from the original LEP1 kit.

### Taxon sampling

To build a backbone phylogeny of the Bombycoidea, we sampled all major lineages (i.e., families, subfamilies, and tribes), with the exception of three rare, species-poor groups: subfamily Munychryiinae (Anthelidae) and tribes Sataspedini and Monardini (Sphingidae), for which representative samples were unavailable for DNA sequencing. Sampled lineages were chosen because: 1) they were appropriate representatives of the taxonomic group needed for the analysis (i.e., good morphological and evolutionary representative of a tribe); and 2) they were accessible for use in phylogenomics. In total, 115 ingroup Bombycoidea species from 97 genera were included in the phylogenetic analysis, as well as two Lasiocampidae outgroups – the sister lineage to the bombycoids (Supp. Table 2).

Specimens were obtained from fieldwork, historically preserved dry collections (Supp. Table 1), and molecular tissue collections. Field-collected specimens were stored in ≥95% ethanol, RNAlater, or papered and dried with silica gel. Genomic DNA was extracted using OmniPrep Genomic DNA Extraction Kits (G-Biosciences, St. Louis, MO, USA) and DNeasy Blood and Tissue Kits (Qiagen, Valencia, CA, USA). DNA concentration was evaluated through agarose gel electrophoresis and fluorometry using a Qubit 2.0 (Invitrogen, Thermo Fisher Scientific, Carlsbad, CA, USA). Library preparation, hybridization enrichment, and Illumina HiSeq 2500 sequencing (PE100) was carried out at RAPiD Genomics (Gainesville, FL, USA). Specimen wing vouchering and tissue storage methods follow Cho et al. [52]. All DNA extracts and specimens preserved in ethanol, RNAlater, or those freshly papered were stored at −80°C at the Florida Museum of Natural History, McGuire Center of Lepidoptera and Biodiversity (MGCL).

### DNA extraction protocol for museum specimens

We evaluated the efficacy of obtaining DNA from historical museum specimens because there is great interest in understanding the feasibility of this approach for use in phylogenetics and systematics ([27], [53], [54], [55]). The Lepidoptera specimens evaluated herein, were “field-pinned” (never rehydrated), “papered” (stored in an envelope and kept dry since collected, thus not rehydrated or pinned), and “traditionally-pinned” specimens (dried, rehydrated, and subsequently pinned) – the historically most common method of Lepidoptera specimen storage. Collecting dates ranged from 1987 to 2017 (Supp. Table 2). Samples were not initially intended to be preserved for molecular sequencing and information about potential contaminants and/or extraction inhibitors (i.e. fumigation compounds, other chemicals) was unavailable. In some cases there was little soft-tissue to extract from within the abdomens, having degraded over time. Being that these samples were not kept in molecular-grade conditions, many were contaminated with fungal and bacterial growth, which could be identified visually by the spores left on the bodies or by the smell of decay. These factors along with the amount of sclerotized tissue present and the fact that the abdomens used needed to stay intact (not homogenized) for dissecting purposes, made extracting good quality genomic DNA challenging.

Our extraction method, detailed in the Supplemental information, attempts to account for several factors: the amount of degraded tissue, the presence of eggs, the relative fat content, and the overall abdomen size. Many commercial DNA extraction kits on the market (including the Omni Prep kit used in this study) recommend using 10 mg-20 mg of well preserved, soft tissue for the extraction process. Given that the museum specimens used had been desiccated for many years, a number of abdomens had little to no visible internal soft tissue remaining. To digest the remaining material in solution, we increased the ratio of proteinase K to lysis buffer. Companies that produce DNA extractions kits know, as can be seen by their specific extraction kits, the relative fat content of samples is an issue because lipid-rich tissue can interfere with the digestion of the soft tissue as well as change the chemistry of the DNA isolation buffers. Specimens that appeared to be “greasy” or seemed to have an oily film on the abdomen were not used for extraction. The overall size of the abdomen was used to estimate the amount of lysis buffer needed in order to sufficiently submerge the abdomen and reach the available soft tissue. The amount of buffer also reflected the volume of reagents needed for the remainder of the DNA isolation process. Lastly, the modified protocol also allows a user to easily prepare the genitalia for taxonomic or morphological taxonomic work as these structures remain undamaged.

### Bioinformatics

The bioinformatics pipeline of Breinholt et al. [16] was used to clean and assemble raw Illumina reads for each AHE locus. The pipeline uses a probe-baited iterative assembly that extends beyond the probe region, checks for quality and cross contamination due to barcode leakage, removes paralogs, and returns a set of aligned orthologs for each locus and taxon of interest. To accomplish these tasks, the pipeline uses the *Bombyx mori* genome [44], and an AHE reference library, which in this study was the BOM1 reference library.

Loci for phylogenetic analysis were selected by applying a cutoff of ≥40% sampled taxa recovery (i.e., for a locus to be included in the analysis, the locus had to be recovered in at least 40% of the sampled taxa). The pipeline evaluates density and entropy at each site of a nucleotide sequence alignment. We elected to trim with entropy and density cutoffs only in “flanking” regions, allowing the “probe” region (exon) to be converted into amino acid sequences. For a site (outside of the probe region) to remain, that site must pass a 60% density and 1.5 entropy cutoff, rejecting sites that fail these requirements. A higher first value (60) increases the coverage cutoff (e.g., a site is kept if 60% of all taxa are represented at that site). A higher second value (1.5) increases the entropy cutoff (i.e., entropy values represent the amount of saturation at a site); sites with values higher than 1.5 possess higher saturation and are thus deleted). AliView v1.18 [56] was used to translate to amino acids, check for frame shifts, recognize and remove stop codons, and edit sequencing errors or lone/dubious indels. Because flanking sequences are generally non-coding and sites have been deemed homologous (see [16]), these flanking sequences, before and after the probe regions, were separated from the exons, then combined and treated together as an independent partition. Due to the filtering steps in the bioinformatics pipeline (i.e., site orthology, and density and saturation evaluation), the flanking partition can be viewed as a SNP supermatrix, where each site is homologous, but uninformative sites, saturated sites, or sites with large amounts missing data have been removed.

Of the 115 bombycoid and two outgroup specimens, 110 were sequenced directly using AHE target capture sequencing, of which 68 were sequenced using the BOM1 and 42 using the LEP1 kit. Seven specimens had their AHE loci probe regions mined from previously sequenced transcriptomes or the *B. mori* genome (Supp. Table 2). These specimens did not have flanking data because of nature of transcriptome data. All specimens were processed using either the ‘Bmori’ (for LEP1) or ‘BOM1’ (for BOM1) reference libraries. Previous Breinholt et al., [16] a scripts are available in Dryad [45]. Instructions on how to use the pipeline and additional scripts that were not part of Breinholt et al. [16] are provided in Dryad (will update upon acceptance).

### Phylogenetics

Lepidoptera AHE probe sets comprise highly-conserved coding probe regions (i.e., exons) and more variable, generally non-coding flanking regions (e.g., introns or intergenic regions) located on either side of the probe region [16]. We evaluated both nucleotide and amino acid datasets to examine phylogenetic signal and the role that saturation may play in the data (see [19]). With the Breinholt et al. [16] pipeline and scripts [45], three datasets were built for phylogeny inference: 1) AA = an amino acid supermatrix composed of translated probe region loci; 2) Pr+Fl = a probe + flanking supermatrix; and 3) ASTRAL = the individual loci from the Pr+Fl supermatrix, used for individual gene tree inference and then species tree estimation to evaluate the potential effects of deep coalescence. Additional analyses (i.e., probe region-only, as nucleotides) were investigated, but are not reported here because their outcomes did not differ from those reported.

Concatenated supermatrices were assembled using FASconCAT-G v1.02 [57]. Phylogenetic inference was performed in a maximum likelihood (ML) framework using IQ-TREE MPI multicore v1.5.3 [58]. For both nucleotide and amino acid datasets, the ‘-m TEST’ command was used in IQ-TREE to perform a search for the most appropriate model of amino acid or nucleotide substitution. For all inferences, we performed 1000 random addition sequence (RAS) replicates, and 1000 replicates each for both ultrafast bootstraps (UFBS) (‘–bb’ command) and SH-aLRT tests (‘-alrt’ command). The SH-like approximate likelihood ratio test (SH-aLRT) estimates branch support values that have been shown to be as conservative as the commonly used non-parametric bootstrap values [59]. SH-aLRT and bootstrap values tend to agree for data sets with strong phylogenetic signal (i.e., datasets with loci that are sufficiently large in number of bases, and tips that share sufficient divergence between sequences). Disagreements in branch support are thought to arise as a consequence of small sample size, insufficient data, or saturated divergence levels (see [60]). We classified nodes as “robust” if they were recovered with support values of UFBS ≥ 95 and SH-aLRT ≥ 80 ([59], [60]).

Because concatenation can be misleading when there are high levels of incomplete lineage sorting or deep coalescence [61], we assessed the impact of potential gene-tree discordance ([62], [63], [64]) by inferring a phylogeny for each individual locus, using IQ-TREE under the same parameters as above (i.e., probe and single partition of flanking loci were modeled by site). Species tree estimation was performed in ASTRAL-III [65]. ASTRAL is a computationally efficient and statistically consistent (under the multi-species coalescent) nonparametric method that takes input gene trees and estimates a highly accurate species tree, even when there is a high level of incomplete lineage sorting (or deep coalescence) [66]. The use of ASTRAL is also an informative “data exploration” exercise with phylogenomic datasets, providing valuable information regarding the level of general tree discordance across your set of gene trees, and the potential presence of incomplete lineage sorting/deep coalescence that should be investigated further. To evaluate node support on the species tree, we used the ASTRAL support values (ASV) – local posterior probabilities that are more precise than evaluating bootstrap values across a set of input trees [67]. ASTRAL support values were determined to be “robust” if nodes were recovered with local posterior probabilities ≥ 0.95. All pipeline steps and phylogenomic analyses were conducted on the University of Florida HiPerGator HPC (http://www.hpc.ufl.edu/). All alignment FASTA files, loci information, partition files, tree files, and other essential data files used for phylogenetic inference are available as supplementary materials on Dryad (will update upon acceptance).

### Rogue taxon & outlier locus analyses

We investigated whether our molecular data included rogue taxa or outlier loci that were potentially influencing our phylogenetic results. A rogue taxon analysis was carried out using the online version of RogueNaRok ([68], [69]), http://rnr.h-its.org/, on the 1000 ultrafast bootstrap trees and the consensus tree from the Pr+Fl supermatrix. Outlier taxa and loci analyses were carried out using Phylo-MCOA v.1.4 (PMCoA) [70] in R (on the 650 gene trees used in the ASTRAL analysis). No rogue taxa or loci were found, and therefore we did not prune any taxa or loci from subsequent analyses.

### Diversification rate analyses

As an initial investigation into why some bombycoid lineages are more diverse than others, we examined and quantified how diversification rates (the interplay between speciation and extinction) have changed over time. Simply calculating species diversity per clade and assuming extant diversity is a true indicator of increases in diversification rate could produce significant biases in one’s interpretations due to some charismatic lineages receiving more taxonomic effort than their “boring” sister lineages (see [71], [72], [73], [74]). We therefore applied BAMM [75] and ‘BAMMtools’ [76] to infer the number and location of macroevolutionary rate shifts across our phylogeny, and visualize the 95% credible set of shift configurations.

The fossil record of Lepidoptera, especially of the Bombycoidea, is poor [77]. Prior lepidopteran studies that have included fossils in dating analyses have been scrutinized for incorrect fossil identification or placement on phylogeny [78]. Because Lepidoptera fossils are limited and characters are difficult to discern, we decided not to conduct a dating analysis for this study. Instead, the ML best tree was converted into a relative-rate scaled ultrametric tree using the ‘chronopl’ command in the R package ‘ape’ [51]. This approach produces a tree whose branches are scaled to evolutionary rates, not a dated tree, and provides a way to understand evolutionary changes over relative “time” of the group being investigated.

For the first time, quantifiable rates of diversification were calculated for the Bombycoidea. This is important, because whether a taxonomic group possesses more described species than another related lineage, does not mean they have diversified “more”. A larger number of described species could simply be due to the taxonomic effort, a well-known bias in bombycoids, with the Sphingidae and Saturniidae representing the charismatic groups that most bombycoid taxonomists have historically worked. To account for non-random missing speciation events, we quantified the percentage of taxa sampled within each family and incorporated these in the form of branch-specific sampling fractions. Sampling fractions were based on the updated superfamily numbers calculated from Kitching et al. [1]. Informed priors, based on our sampling and phylogeny, were determined using ‘setBAMMpriors’ in BAMMtools. The MCMC chain was run for 100 million generations, sampling every 1000 generations. Convergence diagnostics was assessed using the R package ‘coda’ [79]. The first 20% of runs were discarded as burn-in.

## Results

Our abdomen soaking approach proved relatively consistent with high enough yield of target DNA to proceed with AHE sequencing. Datasets constructed for phylogeny inference contained the following number of loci, sequence length, and model specifications: 1) AA = 579 loci, 48,456 amino acid residues, modeled by locus; 2) Pr+Fl = 649 probe loci + one flanking locus (261,780 bp), each probe locus was modeled by site, flanking data was maintained as a single partition and modeled by site; and 3) ASTRAL = 650 loci, each locus modeled by site. Differences in loci number between the AA dataset and the Pr+Fl dataset are the result of loci being removed from the AA phylogenetic inference due to a lack of variation at the amino acid level for those loci (i.e., the probe regions were highly conserved with no variation across taxa).

Increasing the number of loci and taxon sampling significantly improved our understanding of Bombycoidea relationships. The inferred relationships are generally consistent across the three phylogenetic inferences that we performed (AA, Pr+Fl, ASTRAL), with all major backbone relationships robustly supported, and all bombycoid families *sensu* Zwick [20] and Zwick et al. [13] were recovered as monophyletic. Due to the methodological approach (i.e., the treatment of different data types) and the more biologically realistic and parsimonious explanation of the topology (see Systematics section in Supplemental Materials), our preferred phylogeny is the tree generated from the “probe + flanking” (Pr+Fl) dataset. All family-level placements of genera *sensu* Kitching et al. [1] were supported, with the exception of *Arotros* Schaus, a genus long considered to be an epiine bombycid [80]. *Arotros* is clearly nested within Apatelodidae in our tree (Fig. 1; Supp. Fig. 1) and is hereby transferred to Apatelodidae (see supplemental text for additional details on this taxonomic change). The family-level rearrangements of Zwick [20] and Zwick et al. [13] were also recovered in our phylogenetic results: the placement of taxa formerly classified in the “Lemoniidae” (e.g., Lemaire & Minet [22]) are recovered within Brahmaeidae; the Apatelodidae are distinct from the Bombycidae and Phiditiidae; and the broader concept of Endromidae, which includes taxa formerly placed within the “Mirinidae” and Bombycidae (e.g., Lemaire & Minet [22]), are recovered as monophyletic.

**Figure 1.**
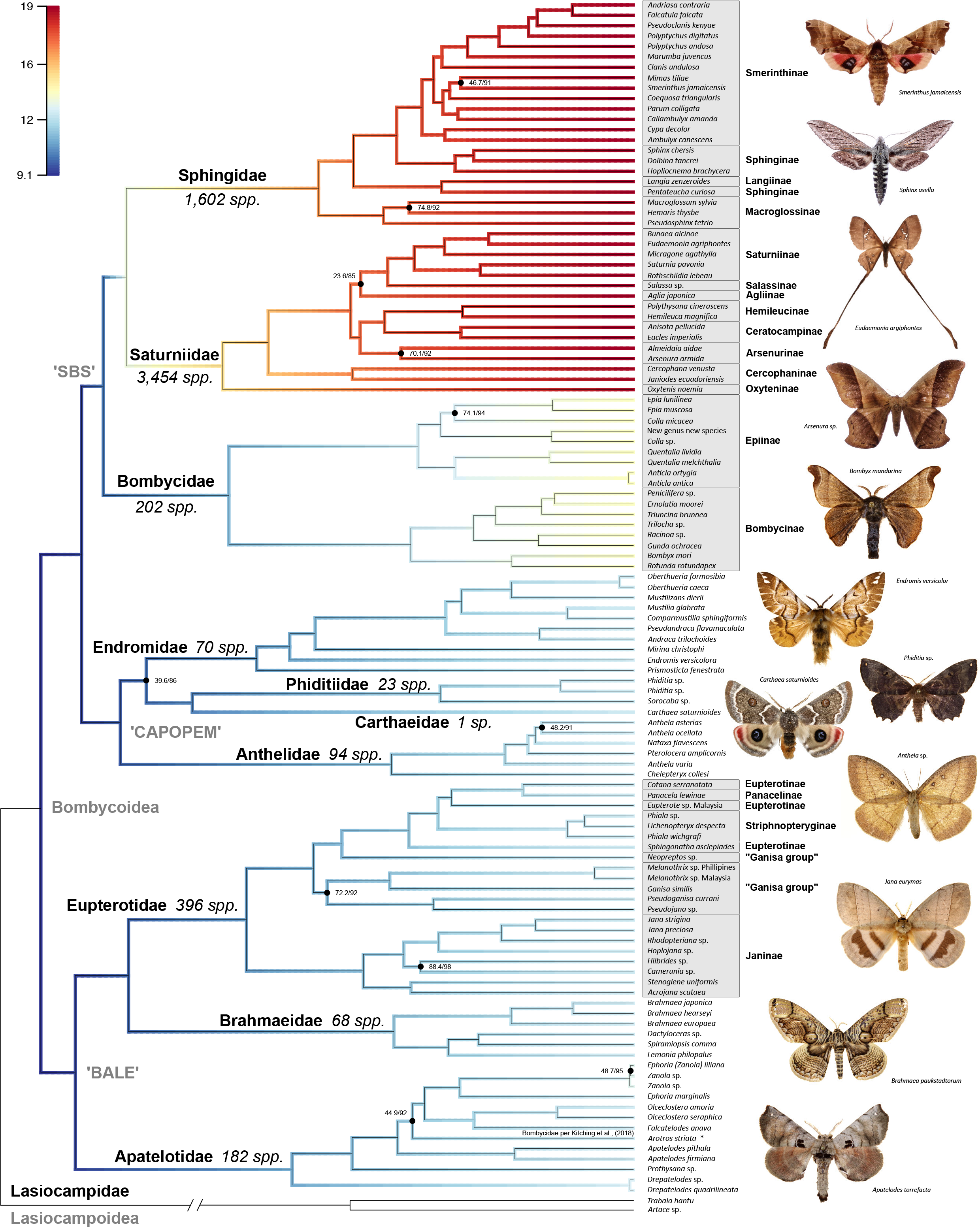
Maximum likelihood tree of Bombycoidea, based on 650 AHE loci. All nodes are supported by ≥95% UFBoot and ≥80% SH-aLRT values unless otherwise noted. Branch color indicates the estimated diversification rate, with warmer colors representing lineages with higher rates. Major taxonomic groups such as families and subfamilies are labeled. Photographs represent species in lineages sampled in the phylogeny. Species diversity, based on Kitching et al. [1], are noted next to families.

We find broad congruence with the major groupings designated by Zwick [20], though internal relationships within these groups did not exactly match previously published trees. Historically, the most problematic familial placement in the superfamily has been the Bombycidae *sensu stricto*. Phylogenetic studies that were based on a handful of gene regions (e.g., [13], [18], [20]), placed this family either as sister to the Saturniidae or to the Sphingidae (reviewed in [19]), albeit without strong support for either. Our study clearly places the Bombycidae as the sister lineage to the Saturniidae + Sphingidae (the ‘SBS’ group – coined by Zwick et al. [13]), as seen in trees from the AA and Pr+Fl datasets, but not in the ASTRAL tree (Supp. Trees 1-3) – an outcome that mirrors traditional Sanger sequencing studies based on few loci, where individual gene trees can lack the phylogenetic signal of supermatrices. The ‘CAPOPEM’ group (Carthaeidae, Anthelidae, Phiditiidae, and Endromidae – coined by Regier et al. [11]) is recovered in all three analyses, although in the ASTRAL inference this group is nested within the clade containing the Sphingidae, Bombycidae and Saturniidae (Supp. Fig. 2).

Interfamilial relationships within the CAPOPEM clade were not robustly supported in Regier et al. [11] or Zwick et al. [13], but our AHE-based trees confidently solidify relationships within this group. Zwick et al. [13] recovered the Old World Endromidae as sister to the Australian Anthelidae + (Australian Carthaeidae + Neotropical Phiditiidae), however, our Pr+Fl analysis supports the Anthelidae as sister to Endromidae + (Carthaeidae + Phiditiidae) instead. Effectively, Anthelidae and Endromidae swap places within CAPOPEM when comparing Zwick et al. [13] and our Pr+Fl phylogeny. Conversely, our AA results illustrate yet a different picture, in which Phiditiidae are sister to Anthelidae + (Carthaeidae + Endromidae), mirroring Regier et al. [11]. This discordance is important to note considering the disjunct geographic distribution of these families. Among these families, only Phiditiidae is found in (and endemic to) the New World, with each of the other families being restricted to the Old World. Lastly, while intrafamilial relationships in the SBS and CAPOPEM groups change depending on which of our datasets are used to infer the phylogeny (Fig. 1, Supp. Figs. 1-3, Supp. Trees 1-3), the ‘BALE’ group (Brahmaeidae, Apatelodidae, and Eupterotidae – coined by Regier et al. [11]) is recovered in all three analyses.

When we investigate intrafamilial relationships, the Bombycidae (excluding *Arotros*, mentioned above; see Supplemental text) possesses distinct and monophyletic subfamilies Bombycinae (Old World) and Epiinae (New World; Fig. 1; Supp. Fig. 1). Within the Eupterotidae, the Striphnopteryginae and Janinae are monophyletic, but others are not (Fig. 1; Supp. Fig. 1). Within the Saturniidae, the Pr+Fl and AA topologies are similar to Regier et al. [31] (Fig. 1 & Supp. Fig. 3), where the Oxyteninae are sister to the rest of the family, followed by the Cercophaninae. Arsenurinae is the sister lineage to the Hemileucinae and the Ceratocampinae, while Salassinae is the sister lineage to the Saturniinae. A major difference in subfamily relationships is the placement of Agliinae, which was either sister to the Salassinae + Saturniinae (Pr+Fl) or to the Arsenurinae (AA, also [13] and [31]). The ASTRAL inference is quite different, placing the Ceratocampinae as the sister lineage to a clade containing the Agliinae, Arsenurinae + Hemileucinae, and a Saturniinae + Salassinae clade (Supp. Fig. 3), albeit with low branch support. Within the Sphingidae, our topologies (both Pr+Fl and AA) are largely congruent, but differ slightly from Kawahara et al. [81], Zwick et al. [13], and Kawahara and Barber [33]. Smerinthinae, which now excludes *Langia* (Kitching et al [1]) is monophyletic and the Sphinginae are polyphyletic (Fig. 1; Supp. Fig. 1). This outcome is robustly supported in the Pr+Fl and ASTRAL trees, although internal relationships slightly differ between the two trees. The placement of the Macroglossinae, Langiinae, and Sphinginae are robustly supported in the AA and Pr+FI tree, but low internal node support obfuscates relationships within Smerinthinae.

Among the other families, a number of subfamilies and genera, as defined by the most current classification of the Bombycoidea [1], are not monophyletic (Fig. 1; Supp. Fig. 1). Within Eupterotidae, the Eupterotinae and the “Ganisa group” are not monophyletic. The Eupterotinae are rendered paraphyletic due to the placements of Panacelinae and Striphnopteryginae. The Striphnopteryginae genus *Phiala* is paraphyletic with respect to *Lichenopteryx*, and the Ganisa group is polyphyletic due to the traditional inclusion of the genus *Neopreptos* [82]. Within the Anthelidae, *Anthela* is paraphyletic due to the placement of *Nataxa* and *Pterolocera*, a finding congruent with Zwick [20]. Within Sphingidae, Sphinginae is polyphyletic due to the placement of *Pentateucha* as sister to the Langiinae, and the Smerinthinae genus *Polyptychus* is paraphyletic.

A number of factors can lead to the appearance of a taxonomic group being more “diverse” than other sister lineages. For example, when simply looking at numbers of described species, taxonomic bias in interest and effort could substantially affect our understanding. This is why it is essential to formally test diversification rates. While there are more described species of Saturniidae and Sphingidae than the rest of the Bombycoidea, that doesn’t mean those two families are necessarily more evolutionarily diverse. For the first time, actual quantifiable rates of diversification were calculated for the Bombycoidea, by evaluating how the interplay between speciation and extinction has changed over relative time. At the family level, a major diversification rate shift occurs along the lineage leading to the Saturniidae and Sphingidae (Fig. 1). The effective sample size of the log-likelihood was 8475.169 and the effective sample size of the number of shift events present in each sample was 18141.4. The 95% credible set of rate shift configurations sampled with BAMM can be seen in Supp. Fig. 4. Unfortunately, due to the poor Bombycoidea fossil record and the limited computational approaches currently available to adequately estimate extinction rates, we are unable to discern whether the differences that exist are due to increases or decreases in speciation, although species diversity numbers, based on the classification of Kitching et al. [1] agree with our diversification results. The number of described species for each of the ten bombycoid families, according to Kitching et al. [1] is: Anthelidae (94 spp.); Apatelotidae (182 spp.); Bombycidae (202 spp.); Brahmaeidae (68 spp.); Carthaeidae (1 sp.); Endromidae (70 spp.); Eupterotidae (396 spp.); Phiditiidae (23 spp.); Saturniidae (3,454 spp.); Sphingidae (1,602 spp.).

## Discussion

Within the past decade, a few studies using molecular sequence data have attempted to resolve the phylogenetic relationships of the Bombycoidea ([11], [13], [20]). None have corroborated the earlier morphology-based hypotheses ([15], [22]). In order to establish a backbone for future evolutionary, comparative, and taxonomic studies, we sampled exemplars from all major lineages in the superfamily and used Anchored Hybrid Enrichment (AHE) phylogenomics to provide a robust phylogeny of the superfamily based on the largest taxonomic and molecular sampling to date.

To achieve dense taxonomic coverage, we included many samples from pinned specimens in natural history collections. We modified previous DNA extraction protocols to increase DNA yield for high-throughput sequencing and modified the AHE probe set developed by Breinholt et al. [16] to more efficiently recover phylogenetically-informative loci within the Bombycoidea. To allow more flexibility in the use of the data (e.g., integration of these samples with “legacy” datasets and CO1 for species identification), we added a selection of traditional Sanger-sequenced loci. To assist future usage of the BOM1 and other AHE probe sets, we expanded the bioinformatic protocol of Breinholt et al. [16] to include step-by-step instructions for running the bioinformatics pipeline.

Our phylogeny largely reinforces the results of earlier molecular works, although phylogenomics finally brings into focus those long problematic relationships, while also identifying important topological where some subfamilies and genera are not recovered as monophyletic. When comparing with trees published in previous works, topological discordances are likely the product of increased locus sampling, which provided significantly more phylogenetic information, as well as morphological homoplasy or convergence that likely obscured the true placement of certain taxonomic groups. For example, our results provide a well-supported placement of the historically troublesome family Bombycidae *sensu stricto*, as the sister lineage to Saturniidae + Sphingidae. An earlier phylogenomic study [19] had provided some evidence in support of this relationship, but its taxon sampling was very limited. Our study highlights how morphological convergence in Bombycoidea has confused our understanding of their evolution. For example, the bombycid genus *Rotunda*, endemic to the Old World, and the apatelodid genus *Arotros*, endemic to the New World, are clearly unrelated (Fig. 1; Supp. Fig. 1). However, they are both astonishingly similar in body size, wing shape, and phenotypic appearance (see Supplemental Information: Systematics). Such results imply that there may be adaptive advantages to evolving particular wing shape and size, as found in other bombycoid lineages such as in the Saturniidae [29]. Furthermore, within the CAPOPEM clade, we see a biogeographical distribution that, based on the AA and ASTRAL topologies, appears to be a more biologically realistic and parsimonious explanation (than the Pr+Fl topology) of the evolutionary history due to the placement of the Neotropical Phiditiidae as sister to the remaining and Old World CAPOPEM families (Supp. Fig. 2). In the future, increased taxon sampling will likely help bring better resolution to both the backbone and internal bombycoid relationships. These types of findings, wherein different dataset types answer different questions, highlight the importance of evaluating phylogenetic data in different ways because phylogenetic signal could be hiding in phylogenomic datasets.

From an evolutionary viewpoint, one of the most interesting results came from the first attempt to quantify the diversification rates across the Bombycoidea, in particular, the dramatic shift in diversification rates leading to the Sphingidae and Saturniidae lineages. While maybe not “surprising”, because of the number of species described, this had never been quantified before in this group. We know that simple numbers of described species does not mean necessarily that they have diversified “more” than other closely related lineages. A number of factors could cause one taxonomic group to appear as if there are more species than another related lineage. Perhaps the most important of these is simply due to the taxonomic effort that has historically been applied to a group, a bias that can certainly be found in the bombycoids, with Sphingidae and Saturniidae representing highly charismatic groups where most bombycoid taxonomists have worked.

These findings are fascinating because the Sphingidae and Saturniidae have contrasting life-history strategies, such as larvae feeding on different kinds of food plants, adults having the ability nectar source [35], and anti-bat strategies ([29], [32], [33], [34]) which are reflected in their morphology (e.g., body sizes and wing shapes) and behavior, including flight speed and maneuverability [35]. In an “arms race”, as has been shown between moths and bats, bat predation selectively removes unfit lineages from the environment, thus increasing the speed of evolution of these surviving lineages. As new traits evolve that can be used to effectively evade the predator, subsequent release from the predatory pressure provides the opportunities to diversify ecologically and behaviorally. Although we did not conduct a divergence time estimation analysis in the present study, the origin of the SBS group has been postulated to be approximately 50 mya [83], with Sphingidae originating soon thereafter [33]. Insectivorous bats are thought to have originated roughly 60 mya, and the diversification of Sphingidae and Saturniidae around that time suggests that the incredible taxonomic diversity within these two families could be in part due to bat-related selection pressures resulting in diverse anti-bat traits.

The hawkmoth and silk moth evolutionary story is certainly more complex than simply reflecting their interactions with bats. Although we postulate that the differences in diversification rates are correlated with bat predation, it is possible that these rate shifts are due to other factors, such as ecological specialization or shifts in host plant usage, both as larvae and adults. Amassing and collating behavioral and ecological datasets for the tips of the Bombycoidea Tree of Life for macroevolutionary comparative investigations is essential to furthering our understanding of this diverse, global superfamily, as well as understanding how bats, ecological traits, and/or biogeographical history may or may not have shaped their diversity. At this time, large trait datasets do not exist for these groups, but are currently being worked on to be included in future trait-dependent diversification analyses with much more complete sampling, at the genus and species level across these families, to truly explore the drivers of Bombycoidea diversity. This research establishes some hypotheses to be further tested when more complete sampling of the Bombycoidea has been completed, and robust trait datasets have been collected.

## Conclusions

Our study finally brings into focus long problematic bombycoid relationships and establishes a backbone for future evolutionary, comparative, and taxonomic studies. Our modified DNA extraction protocol allows Lepidoptera specimens to be readily sequenced from pinned natural history collections, and highlights the flexibility of AHE to generate genomic data from a wide range of museum specimens, both age and preservation method. By allowing researchers to tap into the wealth of biological data residing in natural history collections around the globe, these types of methodologies (e.g., DNA from museum specimens and targeted sequencing capture) will provide the opportunities for us to continually add to our understanding of Lepidoptera and Bombycoidea evolution, as well as refinine our understanding of relationships across the Tree of Life.

## Declarations

### Ethics approval and consent to participate

Not applicable

### Consent for publication

Not applicable

### Availability of data and material

The datasets supporting the results of this article are available in Dryad Data Repository (will update upon acceptance with the unique persistent identifier and hyperlink to datasets in http://format). Supplemental Information includes four figures, one table, plus the full museum specimen DNA extraction protocol, systematic information on Bombycoidea families, bioinformatics pipeline scripts and instructions, alignment FASTA files (nucleotides, amino acids, supermatrices, and individual loci), partition files, tree files, and other essential data files that were inputted or outputted from the phylogenetic inference and analysis. Previous Breinholt et al., [16] bioinformatics scripts are available in Dryad [45].

### Competing interests

The authors declare that they have no competing interests.

### Authors’ Information

ORCID ID - Hamilton: 0000-0001-7263-0755; St Laurent: 0000-0001-6439-5249; Dexter: 0000-0002-5136-6083; Kitching: 0000-0003-4738-5967; Breinholt: 0000-0002-3867-2430; Zwick: 0000-0002-7532-1752; Timmermans: 0000-0002-5024-9053; Barber: 0000-0003-3084-2973; Kawahara: 0000-0002-3724-4610

## Funding

This work was partly supported by NSF grants [IOS-1121807, JRB], [IOS-1121739, AYK], [DBI-1349345, AYK], [DBI-1601369, AYK], [DEB-1557007, AYK], and [PRFB-1612862, CAH]; a NERC grant [NE/P003915/1, IJK]; National Geographic Society grants [CRE 9944-16, JRB]; and postdoctoral support from the Florida Museum of Natural History (to CAH).

## Authors’ Contributions

CAH, AYK, JWB, IJK, MT, AZ designed the study. CAH and JWB analyzed data. CAH, AYK, RAS wrote the paper. CAH, AYK, JRB, KD, IJK, RAS, AZ collected, identified, and processed specimens. All authors discussed results and contributed to the writing of the manuscript.

## Acknowledgements

We thank Seth Bybee, Deborah Glass, Yash Sondhi, Jamie Theobald, and Emmanuel Toussaint. We are indebted to Samantha Epstein, John Heppner, Geena Hill, Nicholas Homziak, Jacqueline Miller, Charles Mitter, Kim Mitter, Lary Reeves, Andrei Sourakov, and Andrew Warren, who provided specimens for this project, contributed help in identifying or organizing specimens, or assisted the project in other ways (e.g., lab work or intellectually). The authors would also like to thank the peer-review process for improving the manuscript. The authors acknowledge University of Florida Research Computing for providing computational resources and support that contributed to the research results reported in this publication. URL: http://researchcomputing.ufl.edu

