## Supplemental Information for "Phylogenomics resolves major relationships and reveals significant diversification rate shifts in the evolution of silk moths and relatives"

All specimens used within this study are housed in: the McGuire Center for Lepidoptera and Biodiversity (MGCL) at the Florida Museum of Natural History, Gainesville, USA (FLMNH); the University of Maryland, College Park, USA (UMD); the Muséum national d'Histoire naturelle in Paris, France (MNHN); and the Australian National Insect Collection in Canberra, Australia (ANIC).

**DNA extraction protocol of dried museum specimens (detailed instructions)**

Prior to tissue sampling, dried (pinned or papered) specimens were assigned MGCL barcodes, photographed, and their labels digitized. Abdomens were then removed using sterile forceps, cleaned with 100% ethanol between each sample, and the remaining specimens were returned to their respective trays within the MGCL collections. Abdomens were placed in 1.5 mL microcentrifuge tubes with the apex of the abdomen in the conical end of the tube. For larger abdomens, 5 mL microcentrifuge tubes or larger were utilized. A solution of proteinase K (Qiagen Cat #19133) and genomic lysis buffer (OmniPrep Genomic DNA Extraction Kit) in a 1:50 ratio was added to each abdomen containing tube, sufficient to cover the abdomen (typically either 300 µL or 500 µL) - similar to the concept used in Hundsdoerfer & Kitching [1]. Ratios of 1:10 and 1:25 were utilized for low quality or rare specimens. Low quality specimens were defined as having little visible tissue inside of the abdomen, mold/fungi growth, or smell of bacterial decay. Samples were incubated overnight (12-18 hours) in a dry air oven at 56°C. Importantly, we also adjusted the ratio depending on the tissue type, i.e., increasing the ratio for particularly large or egg-containing abdomens. Both tissue size and type played a significant role in determining which samples would be best for sequencing. For example, the bigger the abdomen, the more lysis buffer is needed to fully immerse during the digestion incubation step. However, when the abdomen was large and hollow, more proteinase K was used to ensure that protein digestion could take place and not simply cell lysis. Because lipid rich tissue can interfere with the tissue digestion, as well as change the chemistry of the DNA isolation buffers, specimens that appeared "greasy" (i.e., high fat content) were deemed not ideal for this extraction method.

After incubation, samples were spun down briefly using a mini centrifuge to collect any droplets of solution. Abdomens were removed from the extraction buffer using sterile forceps, that had been first cleaned with a 10% bleach solution and then with 100% ethanol between each sample. Abdomens were gently rinsed with ≥95% Ethanol and stored at -20°C in ≥95% Ethanol to await genitalia dissection. This modified protocol allowed DNA extraction without compromising genitalia preparation for future taxonomic or morphological work. For samples containing 300 µL of buffer solution, 100 µL of Chloroform (Sigma Aldrich Cat# C2432-500) was added. For samples utilizing 500 µL of buffer solution, 200 µL of Chloroform was used. Samples were then vortexed and centrifuged for 10 minutes at 14,000 RPM. Supernatant was pipetted off into a new, clean, labeled 1.5 mL microcentrifuge tube. Excess chloroform and sample detritus were discarded into the proper hazardous waste container. For samples containing 300 µL of supernatant, 25 µL of DNA stripping buffer (OmniPrep Kit) was added. For samples consisting of 500 µL of supernatant, 50 µL of DNA stripping buffer was used. Samples were then vortexed and spun down using a mini microcentrifuge, followed by incubation in the dry air oven at 56°C for 10 minutes.

Once incubation was complete, samples were left to cool to room temperature for approximately 10 minutes. For samples containing 300 µL of supernatant, 50 µL of precipitation buffer (OmniPrep Kit) was added. For samples with 500 µL of supernatant 100 µL of precipitation buffer was used. During this step, all samples also received 5 µL of mussel glycogen (OmniPrep Kit). Samples were then vortexed to mix and centrifuged for 20 minutes at 14,000 RPM. The supernatant was pipetted off into a new, sterile, labelled 1.5 mL microcentrifuge tube, leaving a salt pellet in the original tube that was discarded. For samples containing 300 µL of supernatant, 250 µL Isopropanol (Fisher Scientific Cat# BP2618-500) was added to each sample. For samples of 500 µL of supernatant, 500 µL of Isopropanol was used. Samples were vortexed and spun down using a mini centrifuge, then incubated at 4°C for 30 minutes. This step is a "stop step", allowing for the samples to be incubated in the freezer overnight, if necessary.

After incubation, samples were centrifuged for 10 minutes at 14,000 RPM. Excess Isopropanol was removed via pipette into the proper waste container. Once the Isopropanol was removed, 350 µL of 80% Ethanol was added to each sample, followed by the samples being centrifuged for 10 minutes at 14,000 RPM. Ethanol was poured off and samples were spun down using the mini centrifuge so that the remaining ethanol could be pipetted off. Sample tubes were then left open, but covered with a Kimwipe™ for 5-10 minutes to allow the remaining ethanol to evaporate. 50 µL of TE Buffer (OmniPrep Kit) and 0.5 µL of RNAse (OmniPrep Kit) were then added to each sample. Samples were vortexed and spun down using the mini centrifuge and stored at 10°C. DNA quality and quantity were evaluated through agarose gel electrophoresis and fluorometry using a Qubit 2.0 (Invitrogen, Thermo Fisher Scientific) to measure the relative DNA concentrations of each sample. Although yielding high amounts of gDNA, extracts from some museum specimens were too fragmented to be utilized for AHE sequencing. High quality gDNA was determined to be ≥2000bp; ≥1000 bp was deemed “good”; ~300 bp may work, but our experience is that not all of the loci will be recoverable; low quality gDNA is classified as ≤300 bp and will not capture or sequence well. DNA extracts were stored, as above, in the MGCL molecular collection.

**Systematics**

*Sphingidae*

Historically, the Sphingidae have been considered a “highly derived” group (Brock [2], Holloway [3], Lemaire & Minet [4]), which, because of their divergent morphology, have sometimes been assigned to their own monotypic superfamily, Sphingoidea, rather than being included within Bombycoidea, where all recent morphological and molecular phylogenetic studies have now placed them. However, their position within the superfamily, with regard to families Saturniidae and Bombycidae (in the current sense, see below), has been uncertain. The current study confirms a sister relationship between Sphingidae and Saturniidae, with Bombycidae as their sister, in the SBS group.

However, the relationships among the sphingid genera and their assignment to suprageneric taxa have yet to be fully elucidated. The history of the classification of Sphingidae was summarized by Kitching & Cadiou [5], who then provided a slightly updated classification in which they recognized three subfamilies: Smerinthinae, containing three tribes, Smerinthini, Sphingulini, and Ambulycini; Sphinginae, with two tribes, Sphingini and Acherontiini; and Macroglossinae, with three tribes, Dilophonotini, Philampelini, and Macroglossini. The first of these was then divided into two subtribes, Dilophonotina and Hemarina, and the last also into two subtribes, Macroglossina and Choerocampina. Genera in each tribe and subtribe were separated into unnamed subgroups, some of which were indicated as possibly being monophyletic pending further study. Changes from the previous classification were essentially limited to moving Sphingulini from Sphinginae to Smerinthinae (based on larval similarities) and adjusting the placement of some genera.

The classification of Kitching & Cadiou [5] was, however, based on intuitive reasoning rather than phylogenetic analysis. The first phylogenetic analysis was that of Regier et al. [6], based on two nuclear gene sequences. Although the sampling was very limited (14 genera), they recovered strong support for a sister relationship between Smerinthinae and Sphinginae and evidence that the macroglossine tribe Dilophonotini was paraphyletic relative to Philampelini + Macroglossini. Kawahara et al. [7] expanded the sampling to five genes and 50 genera and found numerous novel relationship patterns. Again, Smerinthinae and Sphinginae were recovered as sisters, but only if *Langia* Moore was excluded from the former; this genus was found to be sister to Smerinthinae + Sphinginae. The smerinthine tribe Ambulycini was also monophyletic within a paraphyletic Smerinthini. Tribe Sphingulini was placed with a monophyletic Sphinginae (contra [5] but agreeing with pre-2000 classifications) though paraphyletic, with the Australian genera branching off first, followed by the Asian taxa. Tribe Acherontiini was also recovered as monophyletic, but nested deep within the Sphinginae.

Subfamily Macroglossinae was also recovered as monophyletic. However, subtribe Hemarina was placed as sister to all remaining macroglossines, while subtribe Philampelina, together with a group of macroglossine genera centered on *Proserpinus* Hübner, was nested deep within one of two large clades that comprised tribe Dilophonotina. With the remainder of tribe Macroglossini, subtribe Choercampina was recovered as monophyletic in a paraphyletic subtribe Macroglossina. There were also numerous well supported groups within Smerinthini, Sphingini, Dilophonotina and Macroglossini that suggested further tribal divisions within these groups were warranted. A greatly expanded sampling by Kawahara & Barber [8] found essentially the same pattern of relationships, although subtribe Hemarina was now placed as sister to Macroglossina.

Zolotuhin & Ryabov [9] erected the tribe Sataspedini to accommodate the morphologically divergent and hymenopteran mimetic smerinthine genus *Sataspes* Moore, to which Kitching et al. [12] added the African genus and *Afrosataspes* Basquin & Cadiou. Most recently, Haxaire & Minet [10] proposed two new tribes in Sphinginae, the Pentateuchini and Monardini, to accommodate respectively the genera *Pentateucha* Swinhoe and *Monarda* Druce, and, based on intuitive reasoning, proposed that their relationships within the subfamily were: Pentateuchini + (Monardini + (*Hopliocnema* genus-group + (Sphingulini + Sphingini))).

A new formal higher taxonomy for Sphingidae based on these results and previously implemented in part in the online *Sphingidae Taxonomic Inventory* ([11]; accessed on 5 November, 2018) was published by Kitching et al. [12], in which three subfamilies, Langiinae, Smerinthinae and Sphinginae were recognized. Tribal and subtribal names were resurrected for many of the clades recovered by Kawahara et al. [7] and informal genus-groups proposed where no prior family-rank names were available.

The present study includes only 22 genera of Sphingidae but has nevertheless revealed several previously unknown relationships. The most unexpected was a strongly supported sister paring between *Langia* and *Pentateucha*. This suggests that Pentateuchini should be included as a tribe in Langiinae not Sphinginae. Additionally, Sphinginae could be rendered monophyletic by transferring tribe Monardini from Sphinginae to Langiinae. Several novel relationships are suggested within Smerinthinae, including sister pairs comprising the *Cypa* genus-group + Ambulycini, *Parum* Rothschild & Jordan + *Callambulyx* Rothschild & Jordan (both currently considered unplaced within Smerinthini) and *Coequosa* Walker + (Mimatini + Smerinthini). The inclusion of *Andriasa* Walker in the *Polyptychus* Hübner genus-group is confirmed; unpublished data on pupal morphology had suggested this was likely. Finally, the paraphyly of *Polyptychus* is also not unexpected, although more representatives of this large genus will need to be studied before a formal subdivision of the genus is made.

*Saturniidae*

The Saturniidae are the phylogenetically most well-studied family of bombycoid moths. Each of the subfamilies and tribes (including those that were previously treated as separate families: Cercophaninae and Oxyteninae) have long been recognized to be monophyletic on the basis of strong morphological evidence ([4], [13], [14]). However, the relationships among these clades were poorly understood and often conflicting, even within single works [15]. More recently, Saturniidae have been the focus of several molecular phylogenetic studies ([16], [17],

[18]), or included in broader Bombycoidea or Lepidoptera phylogenetic studies with strong taxon sampling ([15], [19]). Unsurprisingly, the results of these studies have not found any novel subfamily or tribal groups, but they have deepened our understanding of the relationships among these taxa. All recent major molecular phylogenies that included Oxyteninae and Cercophaninae have found these two subfamilies to be sister to the remaining Saturniidae lineages, with Oxyteninae being sister to Cercophaninae + the remainder of the family. However, the relationships among the remaining subfamilies, Agliinae, Arsenurinae, Ceratocampinae, Hemileucinae, Salassinae, and Saturniinae, are less clear.

The present results corroborate the position and relationships of the Cercophaninae and Oxyteninae as sister to the rest of the Saturniidae lineages, the long recognized New World clade comprising Arsenurinae + (Ceratocampinae + Hemileucinae), and the similarly well-established Old World Salassinae + Saturniinae clade. However, a crucial difference from all other molecular phylogenies published thus far is a possible bifurcation between the New World Arsenurinae + (Hemileucinae + Ceratocampinae) and the predominantly Old World Agliinae + (Salassinae + Saturniinae). Previous studies had placed Agliinae either as sister to Arsenurinae + (Ceratocampinae + Hemileucinae) ([16], [17], [19]), a pattern supported by our Pr+Fl analysis; or in a sister relationship with Arsenurinae based on larval morphology (shared possession of tree-like scoli, which could be interpreted as symplesiomorphic). However, our AA analysis unexpectedly suggested that Agliinae were sister to Salassinae + Saturniinae, thereby increasing ambiguity in our understanding of the relationships of this small Old World subfamily.

Yet the most problematic suprageneric relationships are in Saturniinae, due to the historically uncertain composition of and relationships among the African subgroups: “Bunaeinae/-ini”, “Microgoninae/-ini” and “Urotini”. Regier et al. [16] showed a monophyletic Bunaeini and a paraphyletic Urotini. Barber et al. [17] included representatives of all the major Saturniinae lineages but failed to resolve their relationships fully. Rubin and Hamilton et al. [18] densely sampled Saturniinae, using hundreds of AHE loci, to provide the clearest picture so far of the relationships within this subfamily. These authors, like [16] and [17], did not recognize “Bunaeinae” as a subfamily, but instead included it as a tribe within subfamily Saturniinae, together with tribes Saturniini, Attacini, and the less clearly delimited Urotini and Micragonini. Although the monophyly of and sister relationship between Attacini and Saturniini is supported by all these studies, as well as our own, the relationships among the other predominantly African tribes remain highly contentious. Our sampling of Saturniinae was not dense enough to disambiguate the relationships within this subfamily fully. The recently published results of Rubin and Hamilton et al. [18] did provide further insights, but unfortunately they did not discuss the systematics of the subfamily, despite offering the best published dataset so far. Their study showed that the genus *Usta* Wallengren, previously considered a member of tribe Urotini [12], is actually sister to the tribe Micragonini. The remaining Urotini genera form a paraphyletic grade, nested within a monophyletic tribe Bunaeini.

Lastly, our results provide strong support for the sister-group relationship of Saturniidae and Sphingidae, corroborating an unpublished synapomorphy in adult antennal morphology (A. Zwick, pers. obs.).

*Apatelodidae/Bombycidae/Endromidae/Phiditiidae*

Historically, most genera that are now dispersed among the four families Bombycidae, Apatelodidae, Phiditiidae, and Endromidae were consolidated into a broader concept of Bombycidae ([4], [14]). The exceptions were *Endromis* Ochsenheimer and *Dalailama* Staudinger, which comprised Endromidae, and *Mirina* Staudinger, the sole representative of Mirinidae. This view changed radically with the advent of molecular systematics [19], when the genera *Oberthueria* Kirby ("Oberthueriinae") and *Prismosticta* Butler ("Prismostictinae"), together with *Mirina*, were transferred into a broader concept of Endromidae. At the same time, Epiinae were upgraded from tribal to subfamily status in Bombycidae, and Apatelodidae and Phiditiidae were accorded family status. Our results strongly support the interfamilial findings of Zwick et al. [19], Wang et al. [20], and Lin et al. [21]. There is a clear reciprocally monophyletic bifurcation within the true Bombycidae, leading to an exclusively Neotropical Epiinae and the entirely Old World Bombycinae. The remaining genera previously placed in Bombycidae *sensu lato* are now unequivocally confirmed to be relatively unrelated to true bombycids, and are included within the CAPOPEM group as the families Apatelodidae, Phiditiidae and Endromidae (including the former Mirinidae and the former bombycid subfamilies "Oberthueriinae" and "Prismostictinae").

Within Endromidae, the branches subtending *Prismosticta*, *Endromis*, *Mirina* and the *Oberthueria* group are all quite long and suggest that a subfamily division of Endromidae into Prismostictinae, Endrominae, Mirininae and Oberthueriinae might be considered. However, several key genera, such as *Dalailama*, *Sesquiluna* Forbes and *Theophoba* Fletcher & Nye, remain to be sequenced and their relationships are unconfirmed. Hence, we refrain for the moment from recognizing subfamilies in Endromidae pending further study.

Following Zwick [22] and Zwick et al. [19], we treat Apatelodidae as a well-supported family, well differentiated from Bombycidae, with which they were long considered closely related ([4], [14], [23]). Our study is the first to include more than two apatelodid genera, which confirms the monophyly of the family and provides a much-needed insight into the intrafamilial relationships of this poorly studied group. All our analyses recover the Apatelodidae genera as a grade within a well-supported monophyletic group, without clearly defined subclades, suggesting subfamilies are not warranted in this family. However, we were unable to sequence *Carnotena* Walker, *Thelosia* Schaus and *Thyrioclostera* Draudt, and denser taxon sampling within Apatelodidae may reveal finer details of the intergeneric relationships.

Our results also show that the only sampled genus currently (*sensu* [12]) misplaced in a bombycoid family is *Arotros* Schaus. This genus has been included in Bombycidae since its original description. However, our molecular phylogenetics results show that this unique monotypic genus is actually deeply nested within Apatelodidae. Interestingly, *Arotros* displays a fascinating degree of convergent evolution in habitus with the bombycine genus *Rotunda*, to which it is unrelated, relatively speaking, within Bombycoidea (both *Arotros* and *Rotunda* are sampled here). Based on this robustly supported placement, we transfer *Arotros* to Apatelodidae and recommend further research into its morphology to reinforce its move from Bombycidae.

*Carthaeidae*

This enigmatic, monotypic Australian family has been difficult to place with molecular systematics. Regier et al. [24] recovered it as sister to either Anthelidae or Endromidae, depending on the analysis, whereas Zwick [22] recovered Carthaeidae in various uncertain topological placements. Zwick et al. [19] was the first to recover a better supported sister relationship between Carthaeidae and the small Neotropical Phiditiidae, which was also the most robustly supported placement in our research (Pr+Fl analysis). This placement however, is still inconclusive due to the recovery of a weakly supported sister relationship between Carthaeidae and Endromidae in our AA analysis.

*Brahmaeidae*

A close association between Brahmaeidae and Lemoniidae had long been postulated ([4], [14], [25], [26]), but it was not until Zwick [22] that the two families were combined based on strong phylogenetic evidence. Our results support this decision with genus *Lemonia* Hübner nested among typical brahmaeid genera. We were unable to sample the second "lemoniine" genus, *Sabalia* Walker, but Zwick et al. [19] found strong support for this genus as sister to *Lemonia*.

Our study is the first to use molecular phylogenetics to address the position of the enigmatic African genus *Spiramiopsis* Hampson, which has been variously included in "Sabaliadae" (= "Lemoniidae"), Eupterotidae, Bombycidae and Brahmaeidae. However, our results fully support its inclusion in the last of these families, as sister to *Dactyloceras* Mell. The only brahmaeid genus not yet sampled for molecular phylogenetics is *Calliprogonos* Mell, and its phylogenetic placement within the family remains unknown.

*Lemonia* and *Sabalia* are superficially quite distinct both from each other and from the large, intricately patterned *Brahmaea* and *Dactyloceras* (and, to a lesser extent, the more simply patterned *Calliprogonos* and *Spiramiopsis*). Thus, the clade comprising *Lemonia* and *Sabalia* could conceivably be treated as a subfamily, Lemoniinae. However, the African *Dactyloceras* and *Spiramiopsis*, which are sister to *Lemonia + Sabalia*, if considering [19], and their collective sister genus, *Brahmaea*, would then also need to be treated as subfamilies (and no name exists for the former). Given that the position of *Sabalia* remains to be confirmed and that of *Calliprogonos* to be determined, we do not at this time formally assign names to clades within Brahmaeidae, though it would surely be of interest to determine potential synapomorphies for the *Lemonia*, *Sabalia*, *Spiramiopsis* and *Dactyloceras* clade relative to *Brahmaea* to see whether a subfamilial arrangement is indeed warranted.

*Eupterotidae*

The current suprageneric classification of Eupterotidae follows Forbes [26], as modified by Oberprieler et al. [27] and Nässig & Oberprieler [28], comprising four subfamilies and one informal genus group. The subfamily Striphnopteryginae was considered to be monophyletic, as was Janinae, provided that the rather divergent genera *Hibrildes* Druce and *Tissanga* Aurivillius, previously included in their own subfamilies [4], were included. The composition of Eupterotinae has varied over time depending on how the tribe Cotanini was treated. Minet [14] excluded the tribe from Eupterotinae, transferring it to subfamily Panacelinae, and was followed in this regard by Lemaire & Minet [4], Holloway et al. [29] and Zwick [22]. In contrast, Oberprieler et al. [27] and Nässig & Oberprieler [28] preferred to retain Cotanini (comprising three genera, *Cotana* Walker, *Melanergon* Bethune-Baker and *Paracydas* Bethune-Baker) in the subfamily Eupterotinae. The monophyly of Eupterotinae, with or without Cotanini, remains to be demonstrated. Panacelinae, if Cotanini are excluded, comprises only the single genus, *Panacela* Walker. Finally, the "*Ganisa*-group" was proposed as an informal group for those genera not assignable to any of the four subfamilies and as such is most probably paraphyletic, or even polyphyletic. However, this classification of Eupterotidae is not underpinned by any objective phylogenetic analysis. The only objective study of Eupterotidae to date, using molecular sequence data, is that of Zwick [22], but it included only four genera: *Hoplojana* Aurivillius (Janinae), *Ebbepterote* Oberprieler, Nässig & Edwards (Striphnopteryginae), *Cotana* and *Panacela*.

The present study includes the largest sampling of eupterotid genera so far (18), from all five major subgroups, and our results show both agreement and conflict with the current classification. They confirm the monophyly of the subfamilies Janinae and Striphnopteryginae, as well as the close relationship between the genera *Cotana* and *Panacela*, thus supporting the transfer of the former genus (and its close relatives) to Panacelinae. Within Striphnopteryginae, the genus *Phiala* Wallengren is found to be paraphyletic relative to *Lichenopteryx* C. & R. Felder. However, this is perhaps not surprising as the subfamily has received little recent taxonomic attention and is probably just a dumping ground for superficially similar plain and nondescript species. If *Cotana* is accepted as a panaceline, then the large subfamily Eupterotinae is represented by only two taxa. An undetermined species of *Eupterote* Hübner is sister to Panacelinae, supporting a previous suggestion by Zwick [22] that the latter should be treated as a tribe (Panacelini) of the former. The position of *Sphingognatha* C. & R. Felder as sister to

Striphnopteryginae + (Eupterotinae + Panacelinae) is somewhat unexpected as the genus has been synonymized with *Eupterote* [3]; although reinstated by Oberprieler et al. [27], on the basis of its different male genital morphology. Finally, and rather unexpectedly, the *Ganisa*-group was recovered as monophyletic, except for *Neopreptos* Draudt, which is placed as sister to the Eupterotinae/Striphnopteryginae/Panacelinae clade. This is one of only two New World eupterotid genera and is nested within a large group of Asian genera, which has very interesting biogeographical implications. Its position on a long branch also suggests that it may deserve its own subfamily (probably also including the other New World genus, *Preptos* Schaus).

Our results show several patterns of relationship that are incongruent with the current classification and suggest that the subfamily structure will need marked revision. However, the present study is still far from comprehensive and many other genera should be sampled and their relationships elucidated using phylogenetic analysis before any formal changes are proposed.

*Anthelidae*

A close association between Anthelidae and Lasiocampidae had long been suggested ([2], [14], [4], [30]) but the first comprehensive morphological and molecular phylogenetic analysis of the family [22] rejected this in favor of a position deep within Bombycoidea, in a clade together with Carthaeidae and Endromidae. Two subsequent studies ([19], [31]) corroborated this grouping, and added Phiditiidae, to form the CAPOPEM group. However, no agreement regarding the relationships among the four CAPOPEM families was achieved: Zwick [22] found (Carthaeidae (Anthelidae + Endromidae)); a summary of the results of Regier et al.

[31] was an unresolved polytomy; and Zwick et al. [19] recovered (Endromidae (Anthelidae (Carthaeidae + Phiditiidae))). To these, the present study adds (Anthelidae (Endromidae (Carthaeidae + Phiditiidae))). However, support for all these phylogenetic hypotheses is weak. More complete sampling of genera may help to resolve this ambiguity, including especially, in the present context, the second anthelid subfamily, Munychryiinae.

The only reasonably comprehensive phylogenetic treatment of Anthelidae genera to date [22] found that the genus *Anthela* Walker was polyphyletic with regard to several of the other sampled antheline genera, a result also found in the present study. However, a further sampling within this genus (including the type species, *Anthela ferruginosa* Walker) will be necessary before a reclassification of the family can be proposed.

**Supplemental Table & Figures**

*Supp. Table 1.*

List of the loci names in the modified Bombycoidea-specific AHE probe set BOM1. The first column lists the name of the locus; the second column explains what the locus was named in the LEP1 kit or the name of the "legacy" or vision-related loci; the third column lists the length (bp) of each locus.

*Supp. Table 2.*

List of the total taxa included in the phylogenetic analyses. Specimen names refer to tip names, and include the specimen accession number and preliminary taxonomy. These specimen names can be used to identify a specimen's specific molecular data in the supplemental data. Other pieces of information include: taxonomy (family, subfamily, tribe, genus, and species – per Kitching et al. [12]); storage method of the tissues; whether the data came from an AHE probe set, transcriptome, or genome; and collecting date.

*Supp. Figure 1.*

Maximum likelihood tree of Bombycoidea, based on 650 AHE loci. All nodes are supported by ≥95% bootstrap values unless otherwise noted. Major taxonomic groups such as families, subfamilies, and tribes are labeled. Red circles at the tips correspond to genera that are not monophyletic. Black boxes around the tips correspond to non-monophyletic subfamilies.

*Supp. Figure 2.*

Interfamilial relationships of the CAPOPEM group. These relationships change depending on the data used (Pr+Fl or AA) to infer the phylogeny and the phylogenetic inference (supermatrix or ASTRAL). Values at the nodes of the Pr+Fl or AA trees indicate SH-aLRT/UFBS support, or ASV for the ASTRAL analysis.

*Supp. Figure 3.*

Interfamilial relationships of the Saturniidae. These relationships change depending on the data used (Pr+Fl or AA) to infer the phylogeny and the phylogenetic inference method (supermatrix or ASTRAL). Values at the nodes of the Pr+Fl or AA trees indicate SH-aLRT/UFBS support, or ASV for the ASTRAL analysis.

*Supp. Figure 4.*

The four most common 95% credible set of rate shift configurations sampled with BAMM. Branch color indicates the estimated diversification rate, with warmer colors representing lineages with higher rates. Major taxonomic groups with shifts are labeled. Photographs correspond to the major lineages (i.e., the SBS group) with diversification shifts in the phylogeny.

*Supp. Tree 1.*

The Pr+Fl phylogeny.

*Supp. Tree 2.*

The AA phylogeny.

*Supp. Tree 3.*

The ASTRAL phylogeny.

This information as well as additional Supplemental Information can also be found on Dryad (will update upon acceptance with the unique persistent identifier and hyperlink to datasets in http:// format).

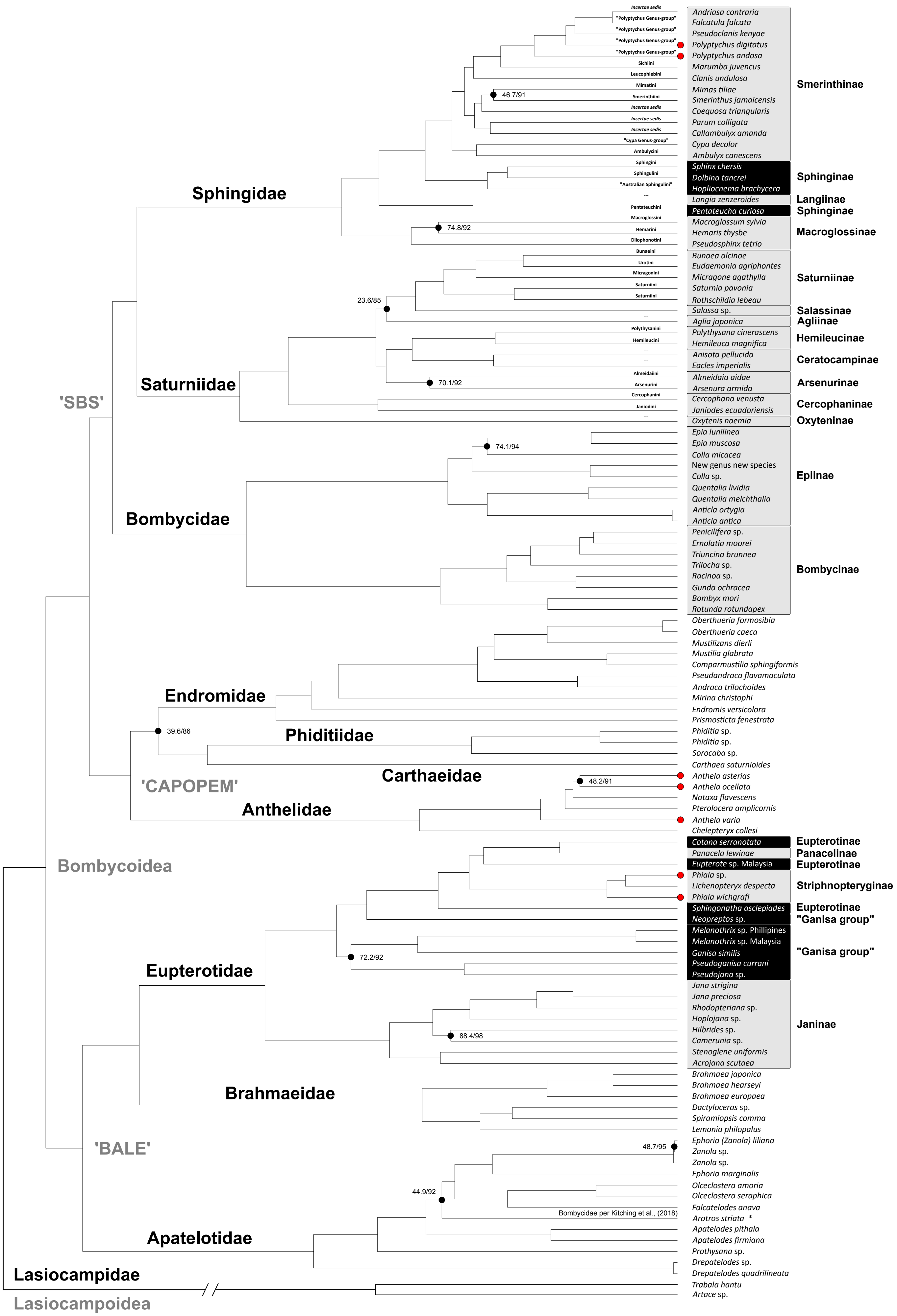

Incertae sedis
"Polyptychus Genus-group"
"Polyptychus Genus-group"
"Polyptychus Genus-group"
"Polyptychus Genus-group"
Sichiini
Leucophlebini
Mimatini
Smerinthini
Incertae sedis
Incertae sedis
Incertae sedis
"Cypa Genus-group"
Ambulycini
Sphingini
Sphingulini
"Australian Sphingulini"
Pentateuchini
Macroglossini
Hemarini
Dilophonotini
Bunaeini
Urotini
Micragonini
Saturniini
Saturniini
Polythysanini
Hemileucini
Almeidaini
Arsenurini
Cercophanini
Janiodini
46.7/91
74.8/92
23.6/85
70.1/92
74.1/94
39.6/86
48.2/91
72.2/92
88.4/98
48.7/95
44.9/92
Andriasa contraria
Falcatula falcata
Pseudoclanis kenyae
Polyptychus digitatus
Polyptychus andosa
Marumba juvencus
Clanis undulosa
Mimas tiliae
Smerinthus jamaicensis
Coequosa triangularis
Parum colligata
Callambulyx amanda
Cypa decolor
Ambulyx canescens
Smerinthinae
Sphinx chersis
Dolbina tancrei
Hopliocnema brachycera
Sphinginae
Langia zenzeroides
Langiinae
Pentateucha curiosa
Sphinginae
Macroglossum sylvia
Hemaris thysbe
Pseudosphinx tetrio
Macroglossinae
Bunaea alcinoe
Eudaemonia agriphontes
Micragone agathylla
Saturnia pavonia
Rothschildia lebeau
Saturniinae
Salassa sp.
Salassinae
Aglia japonica
Agliinae
Polythysana cinerascens
Hemileuca magnifica
Hemileucinae
Anisota pellucida
Eacles imperialis
Ceratocampinae
Almeidaia aidae
Arsenura armida
Arsenurinae
Cercophana venusta
Janiodes ecuadoriensis
Cercophaninae
Oxytenis naemia
Oxyteninae
Epia lunilinea
Epia muscosa
Colla micacea
New genus new species
Colla sp.
Quentalia lividia
Quentalia melchthalia
Anticla ortygia
Anticla antica
Epiinae
Penicilifera sp.
Ernolatia moorei
Triuncina brunnea
Trilocha sp.
Racinoa sp.
Gunda ochracea
Bombyx mori
Rotunda rotundapex
Bombycinae
Oberthueria formosibia
Oberthueria caeca
Mustilizans dierli
Mustilia glabrata
Comparmustilia sphingiformis
Pseudandraca flavamaculata
Andraca trilochoides
Mirina christophi
Endromis versicolora
Prismosticta fenestrata
Phiditia sp.
Phiditia sp.
Sorocaba sp.
Carthaea saturnioides
Anthela asterias
Anthela ocellata
Nataxa flavescens
Pterolocera amplicornis
Anthela varia
Chelepteryx collesi
Cotana serranotata
Eupterotinae
Panacela lewinae
Panacelinae
Eupterote sp. Malaysia
Eupterotinae
Phiala sp.
Lichenopteryx despecta
Phiala wichgrafi
Striphnopteryginae
Sphingonatha asclepiades
Eupterotinae
Neopreptos sp.
"Ganisa group"
Melanothrix sp. Phillipines
Melanothrix sp. Malaysia
Ganisa similis
Pseudoganisa currani
Pseudojana sp.
"Ganisa group"
Jana strigina
Jana preciosa
Rhodopteriana sp.
Hoplojana sp.
Hilbrides sp.
Camerunia sp.
Stenoglene uniformis
Acrojana scutaea
Janinae
Brahmaea japonica
Brahmaea hearseyi
Brahmaea europaea
Dactyloceras sp.
Spiramiopsis comma
Lemonia philopalus
Ephoria (Zanola) liliana
Zanola sp.
Zanola sp.
Ephoria marginalis
Olceclostera amoria
Olceclostera seraphica
Falcatelodes anava
Arotros striata *
Apatelodes pithala
Apatelodes firmiana
Prothysana sp.
Drepatelodes sp.
Drepatelodes quadrilineata
Trabala hantu
Artace sp.
Bombycidae per Kitching et al., (2018)
Sphingidae
Saturniidae
'SBS'
Bombycidae
Endromidae
Phiditiidae
Carthaeidae
'CAPOPEM'
Anthelidae
Bombycoidea
Eupterotidae
Brahmaeidae
'BALE'
Apatelotidae
Lasiocampidae
Lasiocampoidea

'CAPOPEM'

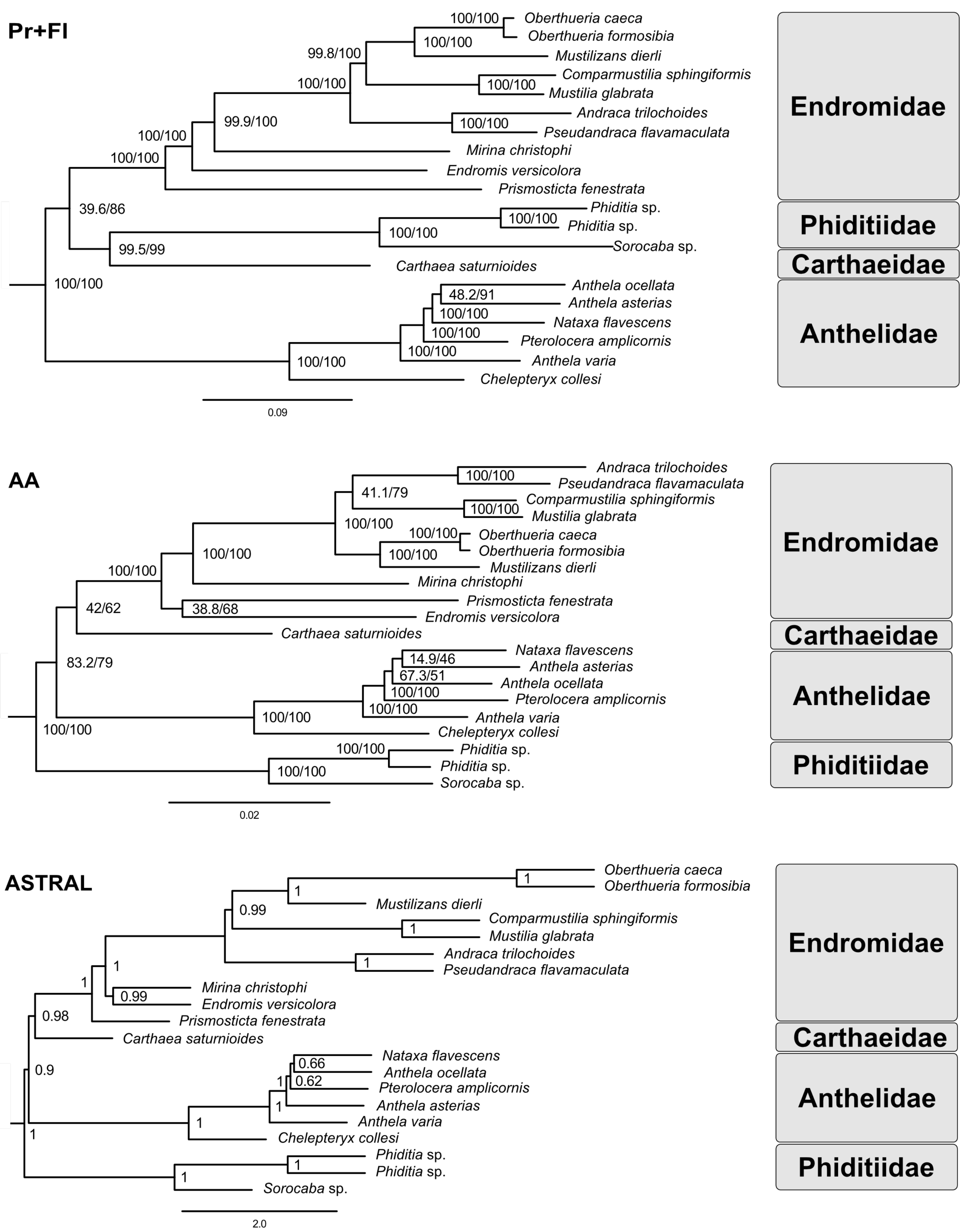

Pr+FI
100/100
Oberthueria caeca
Oberthueria formosibia
100/100
Mustilizans dierli
99.8/100
Comparmustilia sphingiformis
100/100
100/100
Mustilia glabrata
Andraca trilochoides
99.9/100
100/100
Pseudandraca flavamaculata
100/100
Mirina christophi
100/100
Endromis versicolora
Prismosticta fenestrata
39.6/86
Phiditia sp.
100/100
Phiditia sp.
100/100
99.5/99
Sorocaba sp.
Carthaea saturnioides
100/100
Anthela ocellata
48.2/91
Anthela asterias
100/100
Nataxa flavescens
100/100
Pterolocera amplicornis
100/100
Anthela varia
100/100
Chelepteryx collesi
0.09
Endromidae
Phiditiidae
Carthaeidae
Anthelidae
AA
Andraca trilochoides
100/100
Pseudandraca flavamaculata
41.1/79
Comparmustilia sphingiformis
100/100
Mustilia glabrata
100/100
100/100
Oberthueria caeca
Oberthueria formosibia
100/100
100/100
Mustilizans dierli
100/100
Mirina christophi
Prismosticta fenestrata
42/62
38.8/68
Endromis versicolora
Carthaea saturnioides
Nataxa flavescens
14.9/46
Anthela asterias
83.2/79
67.3/51
Anthela ocellata
100/100
Pterolocera amplicornis
100/100
100/100
Anthela varia
Chelepteryx collesi
100/100
100/100
Phiditia sp.
Phiditia sp.
100/100
Sorocaba sp.
0.02
Endromidae
Carthaeidae
Anthelidae
Phiditiidae
ASTRAL
Oberthueria caeca
1
Oberthueria formosibia
1
Mustilizans dierli
0.99
Comparmustilia sphingiformis
1
Mustilia glabrata
Andraca trilochoides
1
1
Pseudandraca flavamaculata
1
Mirina christophi
0.99
Endromis versicolora
0.98
Prismosticta fenestrata
Carthaea saturnioides
Nataxa flavescens
0.66
Anthela ocellata
0.9
1
0.62
Pterolocera amplicornis
1
Anthela asterias
Anthela varia
1
1
Chelepteryx collesi
Phiditia sp.
1
Phiditia sp.
1
Sorocaba sp.
2.0
Endromidae
Carthaeidae
Anthelidae
Phiditiidae

### Saturniidae

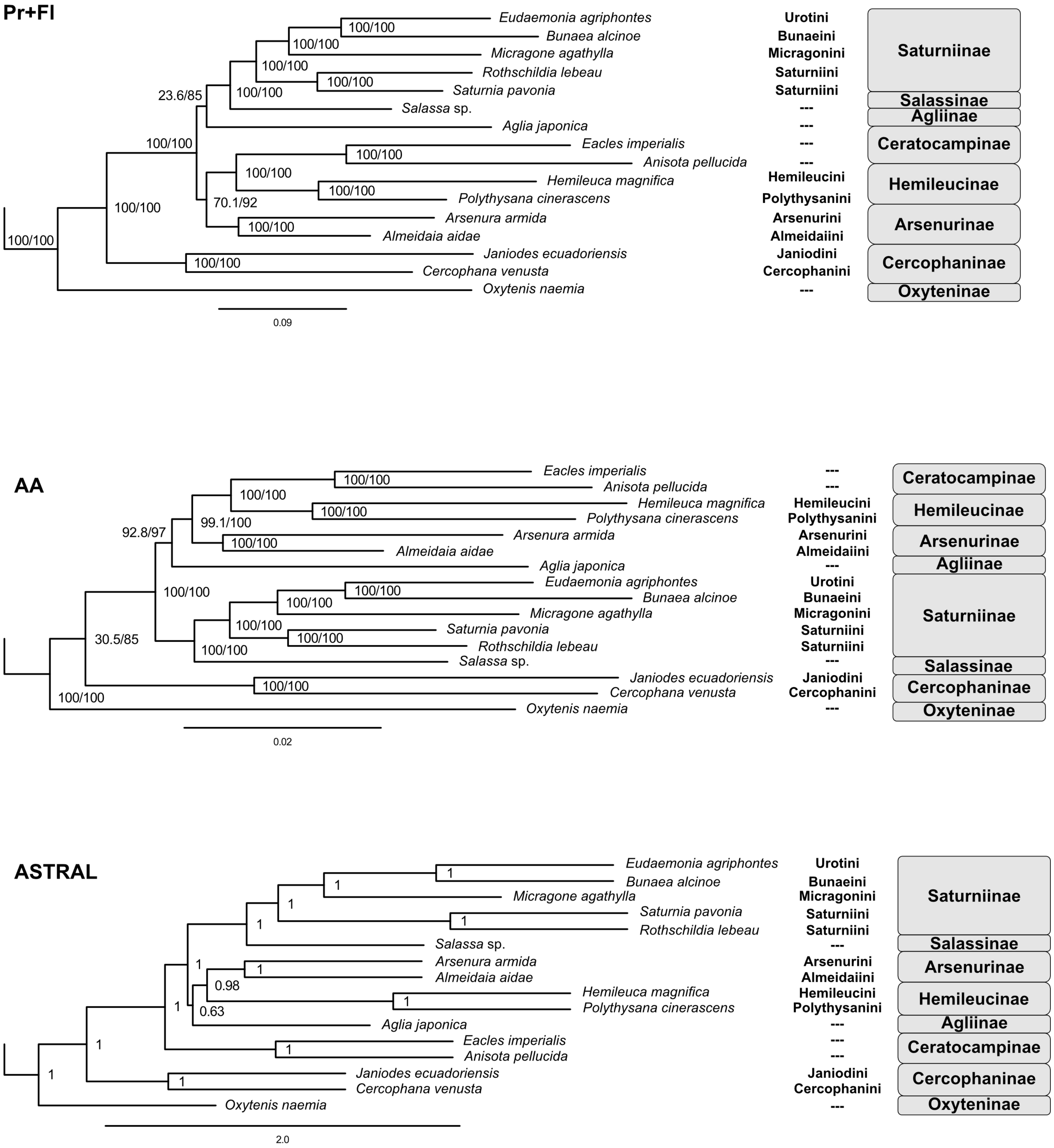

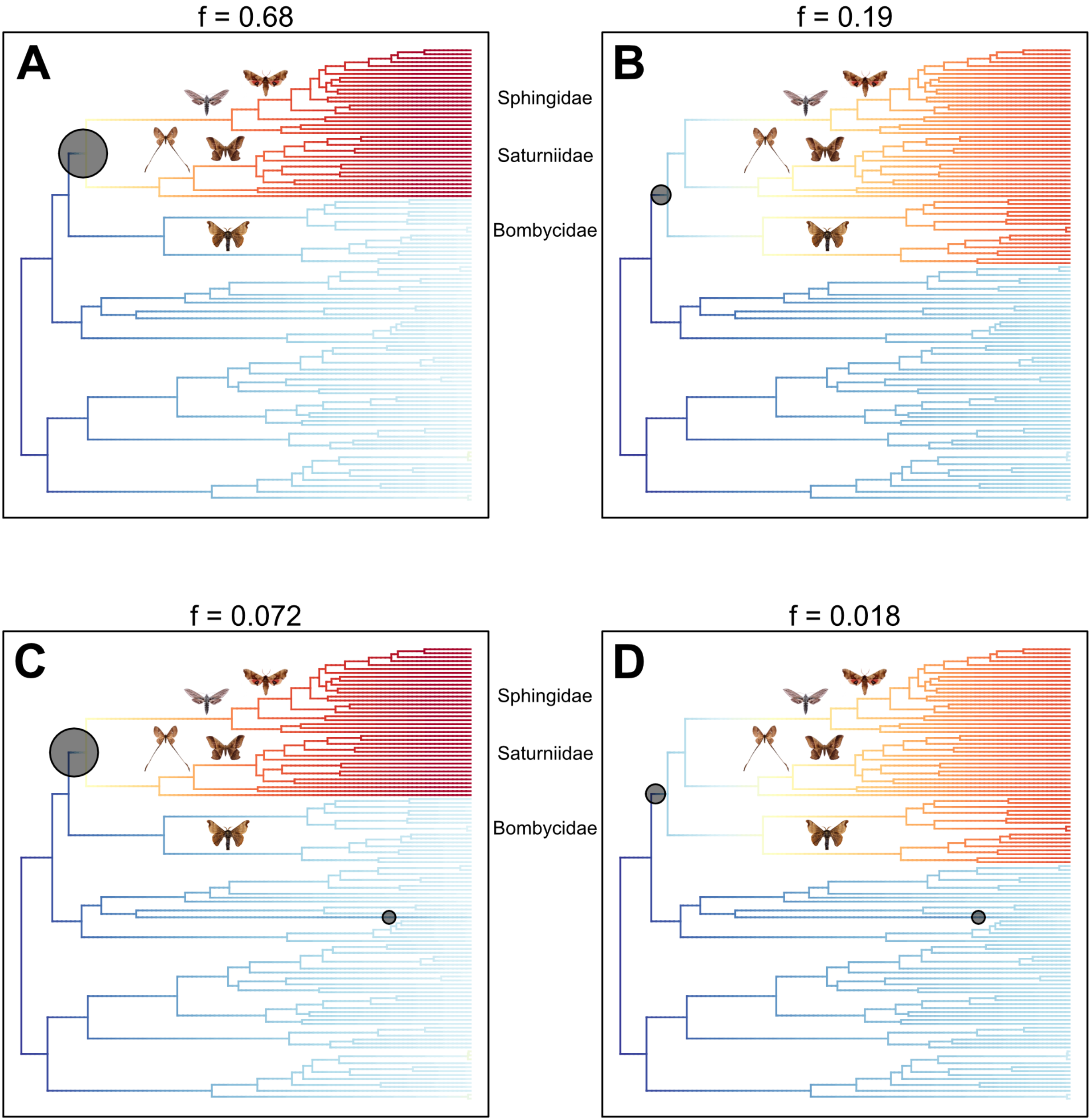
f = 0.68
A
Sphingidae
Saturniidae
Bombycidae
f = 0.19
B
f = 0.072
C
Sphingidae
Saturniidae
Bombycidae
f = 0.018
D

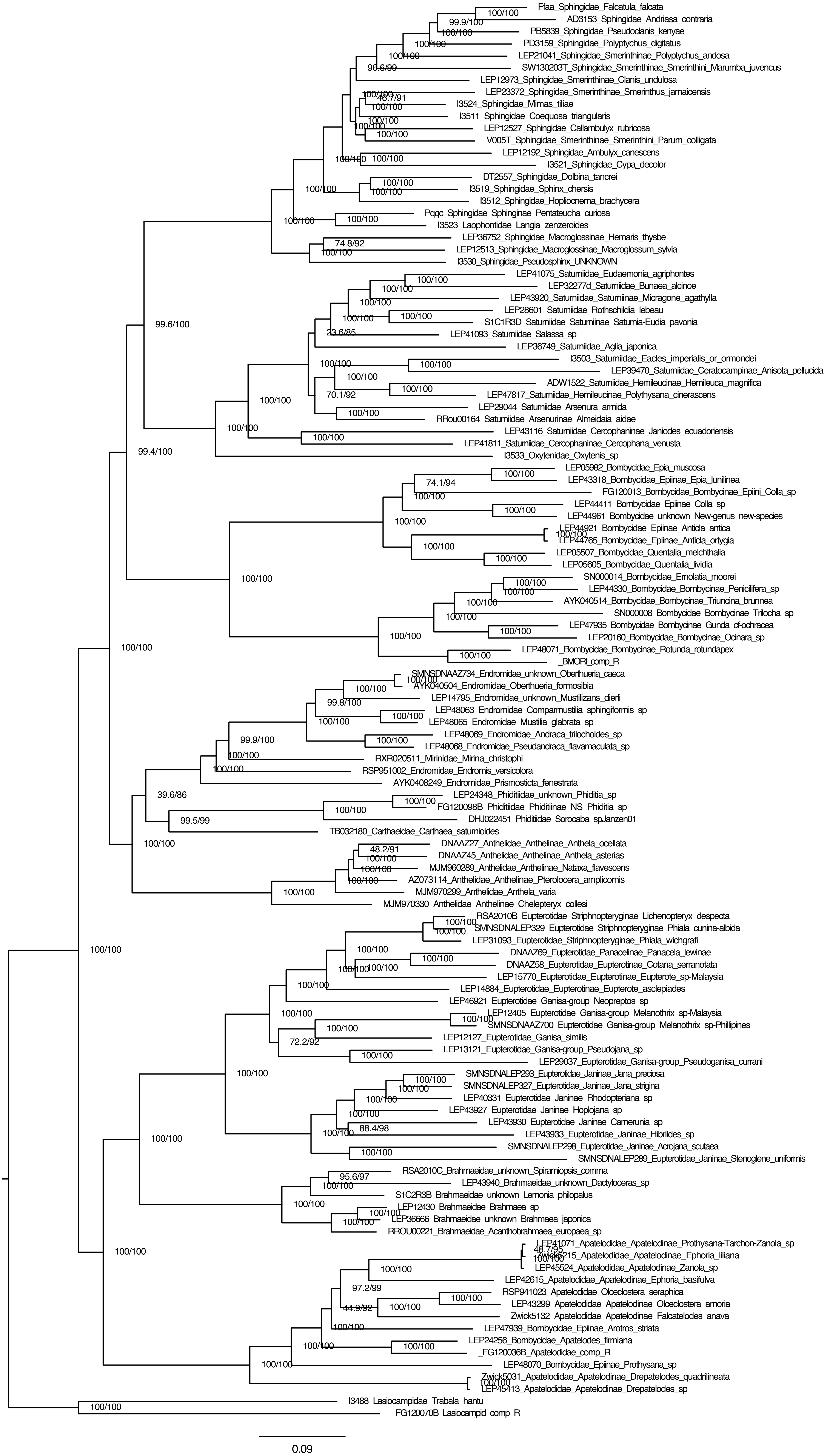
Ffaa_Sphingidae_Falcatula_falcata
AD3153_Sphingidae_Andriasa_contraria
PB5839_Sphingidae_Pseudoclanis_kenyae
PD3159_Sphingidae_Polyptychus_digitatus
LEP21041_Sphingidae_Smerinthinae_Polyptychus_andosa
SW130203T_Sphingidae_Smerinthinae_Smerinthini_Marumba_juvencus
LEP12973_Sphingidae_Smerinthinae_Clanis_undulosa
LEP23372_Sphingidae_Smerinthinae_Smerinthus_jamaicensis
I3524_Sphingidae_Mimas_tiliae
I3511_Sphingidae_Coequosa_triangularis
LEP12527_Sphingidae_Callambulyx_rubricosa
V005T_Sphingidae_Smerinthinae_Smerinthini_Parum_colligata
LEP12192_Sphingidae_Ambulyx_canescens
I3521_Sphingidae_Cypa_decolor
DT2557_Sphingidae_Dolbina_tancrei
I3519_Sphingidae_Sphinx_chersis
I3512_Sphingidae_Hopliocnema_brachycera
Pqqc_Sphingidae_Sphinginae_Pentateucha_curiosa
I3523_Laophontidae_Langia_zenzeroides
LEP36752_Sphingidae_Macroglossinae_Hemaris_thysbe
LEP12513_Sphingidae_Macroglossinae_Macroglossum_sylvia
I3530_Sphingidae_Pseudosphinx_UNKNOWN
LEP41075_Saturniidae_Eudaemonia_agriphontes
LEP32277d_Saturniidae_Bunaea_alcinoe
LEP43920_Saturniidae_Saturniinae_Micragone_agathylla
LEP28601_Saturniidae_Rothschildia_lebeau
S1C1R3D_Saturniidae_Saturniinae_Saturnia-Eudia_pavonia
LEP41093_Saturniidae_Salassa_sp
LEP36749_Saturniidae_Aglia_japonica
I3503_Saturniidae_Eacles_imperialis_or_ormondei
LEP39470_Saturniidae_Ceratocampinae_Anisota_pellucida
ADW1522_Saturniidae_Hemileucinae_Hemileuca_magnifica
LEP47817_Saturniidae_Hemileucinae_Polythysana_cinerascens
LEP29044_Saturniidae_Arsenura_armida
RRou00164_Saturniidae_Arsenurinae_Almeidaia_aidae
LEP43116_Saturniidae_Cercophaninae_Janiodes_ecuadoriensis
LEP41811_Saturniidae_Cercophaninae_Cercophana_venusta
I3533_Oxytenidae_Oxytenis_sp
LEP05982_Bombycidae_Epia_muscosa
LEP43318_Bombycidae_Epiinae_Epia_lunilinea
FG120013_Bombycidae_Bombycinae_Epiini_Colla_sp
LEP44411_Bombycidae_Epiinae_Colla_sp
LEP44961_Bombycidae_unknown_New-genus_new-species
LEP44921_Bombycidae_Epiinae_Anticla_antica
LEP44765_Bombycidae_Epiinae_Anticla_ortygia
LEP05507_Bombycidae_Quentalia_melchthalia
LEP05605_Bombycidae_Quentalia_lividia
SN000014_Bombycidae_Ernolatia_moorei
LEP44330_Bombycidae_Bombycinae_Penicilifera_sp
AYK040514_Bombycidae_Bombycinae_Triuncina_brunnea
SN000008_Bombycidae_Bombycinae_Trilocha_sp
LEP47935_Bombycidae_Bombycinae_Gunda_cf-ochracea
LEP20160_Bombycidae_Bombycinae_Ocinara_sp
LEP48071_Bombycidae_Bombycinae_Rotunda_rotundapex
_BMORI_comp_R
SMNSDNAAZ734_Endromidae_unknown_Oberthueria_caeca
AYK040504_Endromidae_Oberthueria_formosibia
LEP14795_Endromidae_unknown_Mustilizans_dierli
LEP48063_Endromidae_Comparmustilia_sphingiformis_sp
LEP48065_Endromidae_Mustilia_glabrata_sp
LEP48069_Endromidae_Andraca_trilochoides_sp
LEP48068_Endromidae_Pseudandraca_flavamaculata_sp
RXR020511_Mirinidae_Mirina_christophi
RSP951002_Endromidae_Endromis_versicolora
AYK0408249_Endromidae_Prismosticta_fenestrata
LEP24348_Phiditiidae_unknown_Phiditia_sp
FG120098B_Phiditiidae_Phiditiinae_NS_Phiditia_sp
DHJ022451_Phiditiidae_Sorocaba_spJanzen01
TB032180_Carthaeidae_Carthaea_saturnioides
DNAAZ27_Anthelidae_Anthelinae_Anthela_ocellata
DNAAZ45_Anthelidae_Anthelinae_Anthela_asterias
MJM960289_Anthelidae_Anthelinae_Nataxa_flavescens
AZ073114_Anthelidae_Anthelinae_Pterolocera_amplicornis
MJM970299_Anthelidae_Anthela_varia
MJM970330_Anthelidae_Anthelinae_Chelepteryx_collesi
RSA2010B_Eupterotidae_Striphnopteryginae_Lichenopteryx_despecta
SMNSDNALEP329_Eupterotidae_Striphnopteryginae_Phiala_cunina-albida
LEP31093_Eupterotidae_Striphnopteryginae_Phiala_wichgrafi
DNAAZ69_Eupterotidae_Panacelinae_Panacela_lewinae
DNAAZ58_Eupterotidae_Eupterotinae_Cotana_serranotata
LEP15770_Eupterotidae_Eupterotinae_Eupterote_sp-Malaysia
LEP14884_Eupterotidae_Eupterotinae_Eupterote_asclepiades
LEP46921_Eupterotidae_Ganisa-group_Neopreptos_sp
LEP12405_Eupterotidae_Ganisa-group_Melanothrix_sp-Malaysia
SMNSDNAAZ700_Eupterotidae_Ganisa-group_Melanothrix_sp-Phillipines
LEP12127_Eupterotidae_Ganisa_similis
LEP13121_Eupterotidae_Ganisa-group_Pseudojana_sp
LEP29037_Eupterotidae_Ganisa-group_Pseudoganisa_currani
SMNSDNALEP293_Eupterotidae_Janinae_Jana_preciosa
SMNSDNALEP327_Eupterotidae_Janinae_Jana_strigina
LEP40331_Eupterotidae_Janinae_Rhodopteriana_sp
LEP43927_Eupterotidae_Janinae_Hoplojana_sp
LEP43930_Eupterotidae_Janinae_Camerunia_sp
LEP43933_Eupterotidae_Janinae_Hibrildes_sp
SMNSDNALEP298_Eupterotidae_Janinae_Acrojana_scutaea
SMNSDNALEP289_Eupterotidae_Janinae_Stenoglene_uniformis
RSA2010C_Brahmaeidae_unknown_Spiramiopsis_comma
LEP43940_Brahmaeidae_unknown_Dactyloceras_sp
S1C2R3B_Brahmaeidae_unknown_Lemonia_philopalus
LEP12430_Brahmaeidae_Brahmaea_sp
LEP36666_Brahmaeidae_unknown_Brahmaea_japonica
RROU00221_Brahmaeidae_Acanthobrahmaea_europaea_sp
LEP41071_Apatelodidae_Apatelodinae_Prothysana-Tarchon-Zanola_sp
Zwick5215_Apatelodidae_Apatelodinae_Ephoria_liliana
LEP45524_Apatelodidae_Apatelodinae_Zanola_sp
LEP42615_Apatelodidae_Apatelodinae_Ephoria_basifulva
RSP941023_Apatelodidae_Olceclostera_seraphica
LEP43299_Apatelodidae_Apatelodinae_Olceclostera_amoria
Zwick5132_Apatelodidae_Apatelodinae_Falcatelodes_anava
LEP47939_Bombycidae_Epiinae_Arotros_striata
LEP24256_Bombycidae_Apatelodes_firmiana
_FG120036B_Apatelodidae_comp_R
LEP48070_Bombycidae_Epiinae_Prothysana_sp
Zwick5031_Apatelodidae_Apatelodinae_Drepatelodes_quadrilineata
LEP45413_Apatelodidae_Apatelodinae_Drepatelodes_sp
I3488_Lasiocampidae_Trabala_hantu
_FG120070B_Lasiocampid_comp_R
0.09

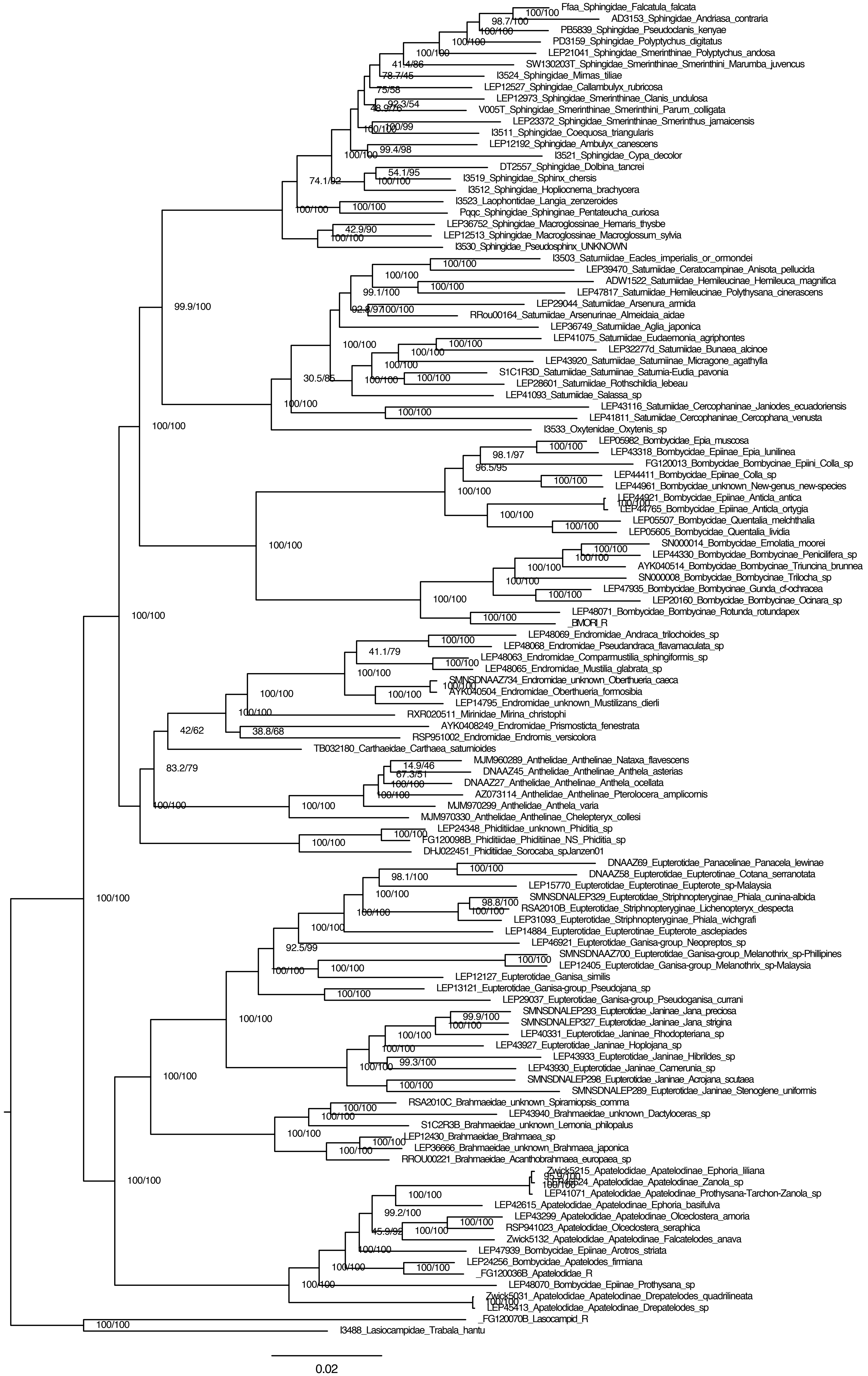
Ffaa_Sphingidae_Falcatula_falcata
AD3153_Sphingidae_Andriasa_contraria
PB5839_Sphingidae_Pseudoclanis_kenyae
PD3159_Sphingidae_Polyptychus_digitatus
LEP21041_Sphingidae_Smerinthinae_Polyptychus_andosa
SW130203T_Sphingidae_Smerinthinae_Smerinthini_Marumba_juvencus
I3524_Sphingidae_Mimas_tiliae
LEP12527_Sphingidae_Callambulyx_rubricosa
LEP12973_Sphingidae_Smerinthinae_Clanis_undulosa
V005T_Sphingidae_Smerinthinae_Smerinthini_Parum_colligata
LEP23372_Sphingidae_Smerinthinae_Smerinthus_jamaicensis
I3511_Sphingidae_Coequosa_triangularis
LEP12192_Sphingidae_Ambulyx_canescens
I3521_Sphingidae_Cypa_decolor
DT2557_Sphingidae_Dolbina_tancrei
I3519_Sphingidae_Sphinx_chersis
I3512_Sphingidae_Hopliocnema_brachycera
I3523_Laophontidae_Langia_zenzeroides
Pqqc_Sphingidae_Sphinginae_Pentateucha_curiosa
LEP36752_Sphingidae_Macroglossinae_Hemaris_thysbe
LEP12513_Sphingidae_Macroglossinae_Macroglossum_sylvia
I3530_Sphingidae_Pseudosphinx_UNKNOWN
I3503_Saturniidae_Eacles_imperialis_or_ormondei
LEP39470_Saturniidae_Ceratocampinae_Anisota_pellucida
ADW1522_Saturniidae_Hemileucinae_Hemileuca_magnifica
LEP47817_Saturniidae_Hemileucinae_Polythysana_cinerascens
LEP29044_Saturniidae_Arsenura_armida
RRou00164_Saturniidae_Arsenurinae_Almeidaia_aidae
LEP36749_Saturniidae_Aglia_japonica
LEP41075_Saturniidae_Eudaemonia_agriphontes
LEP32277d_Saturniidae_Bunaea_alcinoe
LEP43920_Saturniidae_Saturniinae_Micragone_agathylla
S1C1R3D_Saturniidae_Saturniinae_Saturnia-Eudia_pavonia
LEP28601_Saturniidae_Rothschildia_lebeau
LEP41093_Saturniidae_Salassa_sp
LEP43116_Saturniidae_Cercophaninae_Janiodes_ecuadoriensis
LEP41811_Saturniidae_Cercophaninae_Cercophana_venusta
I3533_Oxytenidae_Oxytenis_sp
LEP05982_Bombycidae_Epia_muscosa
LEP43318_Bombycidae_Epiinae_Epia_lunilinea
FG120013_Bombycidae_Bombycinae_Epiini_Colla_sp
LEP44411_Bombycidae_Epiinae_Colla_sp
LEP44961_Bombycidae_unknown_New-genus_new-species
LEP44921_Bombycidae_Epiinae_Anticla_antica
LEP44765_Bombycidae_Epiinae_Anticla_ortygia
LEP05507_Bombycidae_Quentalia_melchthalia
LEP05605_Bombycidae_Quentalia_lividia
SN000014_Bombycidae_Ernolatia_moorei
LEP44330_Bombycidae_Bombycinae_Penicilifera_sp
AYK040514_Bombycidae_Bombycinae_Triuncina_brunnea
SN000008_Bombycidae_Bombycinae_Trilocha_sp
LEP47935_Bombycidae_Bombycinae_Gunda_cf-ochracea
LEP20160_Bombycidae_Bombycinae_Ocinara_sp
LEP48071_Bombycidae_Bombycinae_Rotunda_rotundapex
_BMORI_R
LEP48069_Endromidae_Andraca_trilochoides_sp
LEP48068_Endromidae_Pseudandraca_flavamaculata_sp
LEP48063_Endromidae_Comparmustilia_sphingiformis_sp
LEP48065_Endromidae_Mustilia_glabrata_sp
SMNSDNAAZ734_Endromidae_unknown_Oberthueria_caeca
AYK040504_Endromidae_Oberthueria_formosibia
LEP14795_Endromidae_unknown_Mustilizans_dierli
RXR020511_Mirinidae_Mirina_christophi
AYK0408249_Endromidae_Prismosticta_fenestrata
RSP951002_Endromidae_Endromis_versicolora
TB032180_Carthaeidae_Carthaea_saturnioides
MJM960289_Anthelidae_Anthelinae_Nataxa_flavescens
DNAAZ45_Anthelidae_Anthelinae_Anthela_asterias
DNAAZ27_Anthelidae_Anthelinae_Anthela_ocellata
AZ073114_Anthelidae_Anthelinae_Pterolocera_amplicornis
MJM970299_Anthelidae_Anthela_varia
MJM970330_Anthelidae_Anthelinae_Chelepteryx_collesi
LEP24348_Phiditiidae_unknown_Phiditia_sp
FG120098B_Phiditiidae_Phiditiinae_NS_Phiditia_sp
DHJ022451_Phiditiidae_Sorocaba_spJanzen01
DNAAZ69_Eupterotidae_Panacelinae_Panacela_lewinae
DNAAZ58_Eupterotidae_Eupterotinae_Cotana_serranotata
LEP15770_Eupterotidae_Eupterotinae_Eupterote_sp-Malaysia
SMNSDNALEP329_Eupterotidae_Striphnopteryginae_Phiala_cunina-albida
RSA2010B_Eupterotidae_Striphnopteryginae_Lichenopteryx_despecta
LEP31093_Eupterotidae_Striphnopteryginae_Phiala_wichgrafi
LEP14884_Eupterotidae_Eupterotinae_Eupterote_asclepiades
LEP46921_Eupterotidae_Ganisa-group_Neopreptos_sp
SMNSDNAAZ700_Eupterotidae_Ganisa-group_Melanothrix_sp-Phillipines
LEP12405_Eupterotidae_Ganisa-group_Melanothrix_sp-Malaysia
LEP12127_Eupterotidae_Ganisa_similis
LEP13121_Eupterotidae_Ganisa-group_Pseudojana_sp
LEP29037_Eupterotidae_Ganisa-group_Pseudoganisa_currani
SMNSDNALEP293_Eupterotidae_Janinae_Jana_preciosa
SMNSDNALEP327_Eupterotidae_Janinae_Jana_strigina
LEP40331_Eupterotidae_Janinae_Rhodopteriana_sp
LEP43927_Eupterotidae_Janinae_Hoplojana_sp
LEP43933_Eupterotidae_Janinae_Hibrildes_sp
LEP43930_Eupterotidae_Janinae_Camerunia_sp
SMNSDNALEP298_Eupterotidae_Janinae_Acrojana_scutaea
SMNSDNALEP289_Eupterotidae_Janinae_Stenoglene_uniformis
RSA2010C_Brahmaeidae_unknown_Spiramiopsis_comma
LEP43940_Brahmaeidae_unknown_Dactyloceras_sp
S1C2R3B_Brahmaeidae_unknown_Lemonia_philopalus
LEP12430_Brahmaeidae_Brahmaea_sp
LEP36666_Brahmaeidae_unknown_Brahmaea_japonica
RROU00221_Brahmaeidae_Acanthobrahmaea_europaea_sp
Zwick5215_Apatelodidae_Apatelodinae_Ephoria_liliana
LEP41071_Apatelodidae_Apatelodinae_Prothysana-Tarchon-Zanola_sp
LEP42615_Apatelodidae_Apatelodinae_Ephoria_basifulva
LEP43299_Apatelodidae_Apatelodinae_Olceclostera_amoria
RSP941023_Apatelodidae_Olceclostera_seraphica
Zwick5132_Apatelodidae_Apatelodinae_Falcatelodes_anava
LEP47939_Bombycidae_Epiinae_Arotros_striata
LEP24256_Bombycidae_Apatelodes_firmiana
_FG120036B_Apatelodidae_R
LEP48070_Bombycidae_Epiinae_Prothysana_sp
Zwick5031_Apatelodidae_Apatelodinae_Drepatelodes_quadrilineata
LEP45413_Apatelodidae_Apatelodinae_Drepatelodes_sp
_FG120070B_Lasiocampid_R
I3488_Lasiocampidae_Trabala_hantu
0.02

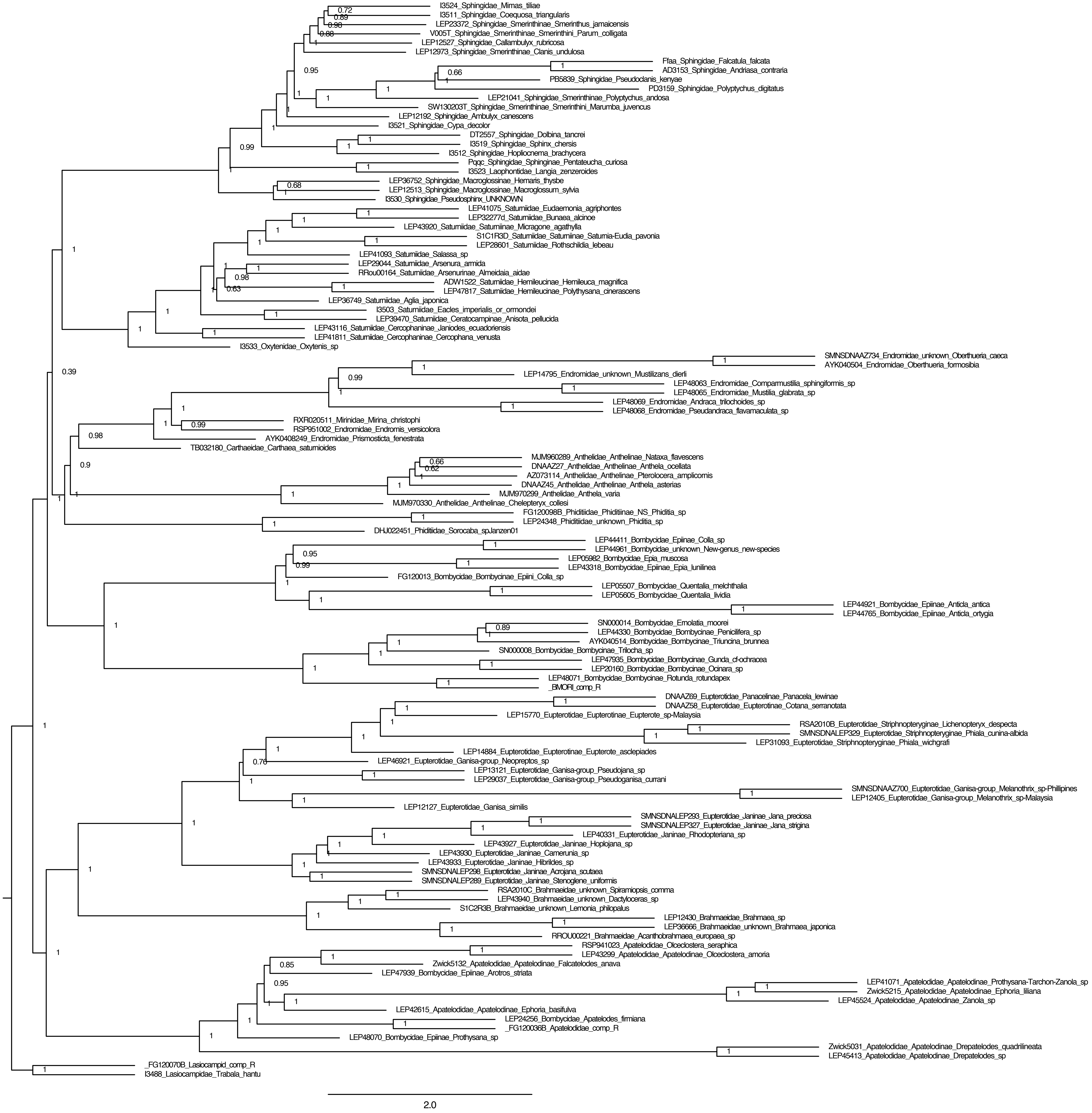
I3524_Sphingidae_Mimas_tiliae
I3511_Sphingidae_Coequosa_triangularis
LEP23372_Sphingidae_Smerinthinae_Smerinthus_jamaicensis
V005T_Sphingidae_Smerinthinae_Smerinthini_Parum_colligata
LEP12527_Sphingidae_Callambulyx_rubricosa
LEP12973_Sphingidae_Smerinthinae_Clanis_undulosa
Ffaa_Sphingidae_Falcatula_falcata
AD3153_Sphingidae_Andriasa_contraria
PB5839_Sphingidae_Pseudoclanis_kenyae
PD3159_Sphingidae_Polyptychus_digitatus
LEP21041_Sphingidae_Smerinthinae_Polyptychus_andosa
SW130203T_Sphingidae_Smerinthinae_Smerinthini_Marumba_juvencus
LEP12192_Sphingidae_Ambulyx_canescens
I3521_Sphingidae_Cypa_decolor
DT2557_Sphingidae_Dolbina_tancrei
I3519_Sphingidae_Sphinx_chersis
I3512_Sphingidae_Hopliocnema_brachycera
Pqqc_Sphingidae_Sphinginae_Pentateucha_curiosa
I3523_Laophontidae_Langia_zenzeroides
LEP36752_Sphingidae_Macroglossinae_Hemaris_thysbe
LEP12513_Sphingidae_Macroglossinae_Macroglossum_sylvia
I3530_Sphingidae_Pseudosphinx_UNKNOWN
LEP41075_Saturniidae_Eudaemonia_agriphontes
LEP32277d_Saturniidae_Bunaea_alcinoe
LEP43920_Saturniidae_Saturniinae_Micragone_agathylla
S1C1R3D_Saturniidae_Saturniinae_Saturnia-Eudia_pavonia
LEP28601_Saturniidae_Rothschildia_lebeau
LEP41093_Saturniidae_Salassa_sp
LEP29044_Saturniidae_Arsenura_armida
RRou00164_Saturniidae_Arsenurinae_Almeidaia_aidae
ADW1522_Saturniidae_Hemileucinae_Hemileuca_magnifica
LEP47817_Saturniidae_Hemileucinae_Polythysana_cinerascens
LEP36749_Saturniidae_Aglia_japonica
I3503_Saturniidae_Eacles_imperialis_or_ormondei
LEP39470_Saturniidae_Ceratocampinae_Anisota_pellucida
LEP43116_Saturniidae_Cercophaninae_Janiodes_ecuadoriensis
LEP41811_Saturniidae_Cercophaninae_Cercophana_venusta
I3533_Oxytenidae_Oxytenis_sp
SMNSDNAAZ734_Endromidae_unknown_Oberthueria_caeca
AYK040504_Endromidae_Oberthueria_formosibia
LEP14795_Endromidae_unknown_Mustilizans_dierli
LEP48063_Endromidae_Comparmustilia_sphingiformis_sp
LEP48065_Endromidae_Mustilia_glabrata_sp
LEP48069_Endromidae_Andraca_trilochoides_sp
LEP48068_Endromidae_Pseudandraca_flavamaculata_sp
RXR020511_Mirinidae_Mirina_christophi
RSP951002_Endromidae_Endromis_versicolora
AYK0408249_Endromidae_Prismosticta_fenestrata
TB032180_Carthaeidae_Carthaea_saturnioides
MJM960289_Anthelidae_Anthelinae_Nataxa_flavescens
DNAAZ27_Anthelidae_Anthelinae_Anthela_ocellata
AZ073114_Anthelidae_Anthelinae_Pterolocera_amplicornis
DNAAZ45_Anthelidae_Anthelinae_Anthela_asterias
MJM970299_Anthelidae_Anthela_varia
MJM970330_Anthelidae_Anthelinae_Chelepteryx_collesi
FG120098B_Phiditiidae_Phiditiinae_NS_Phiditia_sp
LEP24348_Phiditiidae_unknown_Phiditia_sp
DHJ022451_Phiditiidae_Sorocaba_spJanzen01
LEP44411_Bombycidae_Epiinae_Colla_sp
LEP44961_Bombycidae_unknown_New-genus_new-species
LEP05982_Bombycidae_Epia_muscosa
LEP43318_Bombycidae_Epiinae_Epia_lunilinea
FG120013_Bombycidae_Bombycinae_Epiini_Colla_sp
LEP05507_Bombycidae_Quentalia_melchthalia
LEP05605_Bombycidae_Quentalia_lividia
LEP44921_Bombycidae_Epiinae_Anticla_antica
LEP44765_Bombycidae_Epiinae_Anticla_ortygia
SN000014_Bombycidae_Ernolatia_moorei
LEP44330_Bombycidae_Bombycinae_Penicillifera_sp
AYK040514_Bombycidae_Bombycinae_Triuncina_brunnea
SN000008_Bombycidae_Bombycinae_Trilocha_sp
LEP47935_Bombycidae_Bombycinae_Gunda_cf-ochracea
LEP20160_Bombycidae_Bombycinae_Ocinara_sp
LEP48071_Bombycidae_Bombycinae_Rotunda_rotundapex
_BMORI_comp_R
DNAAZ69_Eupterotidae_Panacelinae_Panacela_lewinae
DNAAZ58_Eupterotidae_Eupterotinae_Cotana_serranotata
LEP15770_Eupterotidae_Eupterotinae_Eupterote_sp-Malaysia
RSA2010B_Eupterotidae_Striphnopteryginae_Lichenopteryx_despecta
SMNSDNALEP329_Eupterotidae_Striphnopteryginae_Phiala_cunina-albida
LEP31093_Eupterotidae_Striphnopteryginae_Phiala_wichgrafi
LEP14884_Eupterotidae_Eupterotinae_Eupterote_asclepiades
LEP46921_Eupterotidae_Ganisa-group_Neopreptos_sp
LEP13121_Eupterotidae_Ganisa-group_Pseudojana_sp
LEP29037_Eupterotidae_Ganisa-group_Pseudoganisa_currani
SMNSDNAAZ700_Eupterotidae_Ganisa-group_Melanothrix_sp-Phillipines
LEP12405_Eupterotidae_Ganisa-group_Melanothrix_sp-Malaysia
LEP12127_Eupterotidae_Ganisa_similis
SMNSDNALEP293_Eupterotidae_Janinae_Jana_preciosa
SMNSDNALEP327_Eupterotidae_Janinae_Jana_strigina
LEP40331_Eupterotidae_Janinae_Rhodopteriana_sp
LEP43927_Eupterotidae_Janinae_Hoplojana_sp
LEP43930_Eupterotidae_Janinae_Camerunia_sp
LEP43933_Eupterotidae_Janinae_Hibrildes_sp
SMNSDNALEP298_Eupterotidae_Janinae_Acrojana_scutaea
SMNSDNALEP289_Eupterotidae_Janinae_Stenoglene_uniformis
RSA2010C_Brahmaeidae_unknown_Spiramiopsis_comma
LEP43940_Brahmaeidae_unknown_Dactyloceras_sp
S1C2R3B_Brahmaeidae_unknown_Lemonia_philopalus
LEP12430_Brahmaeidae_Brahmaea_sp
LEP36666_Brahmaeidae_unknown_Brahmaea_japonica
RROU00221_Brahmaeidae_Acanthobrahmaea_europaea_sp
RSP941023_Apatelodidae_Olceclostera_seraphica
LEP43299_Apatelodidae_Apatelodinae_Olceclostera_amoria
Zwick5132_Apatelodidae_Apatelodinae_Falcatelodes_anava
LEP47939_Bombycidae_Epiinae_Arotros_striata
LEP41071_Apatelodidae_Apatelodinae_Prothysana-Tarchon-Zanola_sp
Zwick5215_Apatelodidae_Apatelodinae_Ephoria_liliana
LEP45524_Apatelodidae_Apatelodinae_Zanola_sp
LEP42615_Apatelodidae_Apatelodinae_Ephoria_basifulva
LEP24256_Bombycidae_Apatelodes_firmiana
_FG120036B_Apatelodidae_comp_R
LEP48070_Bombycidae_Epiinae_Prothysana_sp
Zwick5031_Apatelodidae_Apatelodinae_Drepatelodes_quadrilineata
LEP45413_Apatelodidae_Apatelodinae_Drepatelodes_sp
_FG120070B_Lasiocampid_comp_R
I3488_Lasiocampidae_Trabala_hantu
2.0
